# Social isolation of aged mice drives dramatic release of inflammatory lipoxygenase-derived oxylipins

**DOI:** 10.64898/2025.12.15.694286

**Authors:** Mareike Wichmann-Costaganna, Raphaëlle Petit, Julia Lindner, Madlen Haase, Vivien Bachmann, Robert Klaus Hofstetter, Markus Werner, Christiane Frahm, Oliver Werz, Patrick Schädel

**Author notes:** Shared correspondence (Christiane Frahm, Oliver Werz, Patrick Schädel). Address for correspondence (Patrick Schädel): Department of Pharmaceutical/Medicinal Chemistry, Institute of Pharmacy, Friedrich Schiller University Jena, Philosophenweg 14, 07743 Jena, Germany; Address for correspondence (Oliver Werz): Department of Pharmaceutical/Medicinal Chemistry, Institute of Pharmacy, Friedrich Schiller University Jena, Philosophenweg 14, 07743 Jena, Germany; Address for correspondence (Christiane Frahm): Hans Berger Department of Neurology, Jena University Hospital, Am Klinikum 1, 07747 Jena, Germany.

## Abstract

Oxylipins, signalling molecules derived from polyunsaturated fatty acids, act as key mediators controlling inflammatory processes. Ageing fuels the disruption of this network, promoting inflammageing. Social isolation, a common feature of ageing, may contribute to the emergence of pro-inflammatory responses, further aggravating conditions like cognitive decline and frailty. Here, we studied how repeated social isolation impacts inflammation-related oxylipin profiles in seven different organs and serum of aged mice. Additionally, we explored physical exercise as a tool to ameliorate age- and isolation-associated inflammation. Our results show that social isolation induces significant increases in interleukin-1β levels and stimulates a dramatic production of lipoxygenase (LOX)-derived oxylipins in an organ-dependent manner, particularly pronounced in liver, lung, and spleen. Physical exercise failed to mitigate the pro-inflammatory effects induced by social isolation. These effects did not occur in the circulatory system as serum oxylipin levels remained relatively unchanged by isolation. The unexpected and striking elevation of oxylipins across the organs highlights the detrimental effect of social isolation and proposes key roles of oxylipins in stress-related inflammageing.

## Introduction

Ageing is a complex, multifactorial biological and social process, accompanied by the accumulation of deleterious changes often resulting in decreased health, cognitive decline, and overall loss of function and fitness (Harman 2006; Khan, Singer, and Vaughan 2017).

Disruption of homeostasis, caused by impaired immune functions, immunosenescence (Franceschi et al. 2000), and subsequent increase in chronic low-grade inflammation, termed inflammageing (Furman et al. 2019; Fulop et al. 2023), is considered one of the proposed hallmarks of ageing (Lopez-Otin et al. 2023). Inflammatory processes are tightly regulated by a network of mediators, such as cytokines, chemokines, and oxylipins (Franceschi et al. 2018). During ageing, this complex network becomes dysregulated and aggravates inflammatory processes, as oxylipins play vital roles as lipid mediators in facilitating the initiation, perpetuation, and resolution of inflammation (Arnardottir et al. 2014; Chiurchiu, Leuti, and Maccarrone 2018). Oxylipins are produced from polyunsaturated fatty acids (PUFA) including omega-6 (e.g., arachidonic acid (AA, C20:4)) and omega-3 (e.g., docosahexaenoic acid (DHA, C22:6), eicosapentaenoic acid (EPA, C20:5) members. Their mode of action can be both pro- and anti-inflammatory: while pro-inflammatory AA-derived prostaglandins (PG) and leukotrienes (LT) are biosynthesised via the cyclooxygenase (COX) and 5-lipoxygenase (LOX) pathway, anti-inflammatory and pro-resolving oxylipins are mainly omega-3-PUFA-derived and are generated by 5-, 12-, and 15-LOX. The latter oxylipins encompass specialized pro-resolving mediators (SPM) such as protectins, lipoxins (LX), maresins (MaR), resolvins (Rv), as well as their mono-hydroxylated precursors (e.g., 15-hydroxy-eicosapentaenoic acid (15-HEPE), 17-hydroxy-docosahexaenoic acid (17-HDHA)) (Serhan 2014; Leuti et al. 2020). Aberrant oxylipin biosynthesis is a characteristic feature of several age-related diseases, such as cardiovascular disease (Nayeem 2018; Bäck and Hansson 2019), atherosclerosis (Gleim et al. 2012), and neurodegeneration (Whittington, Planel, and Terrando 2017; Chiurchiu et al. 2022). In recent studies, we revealed that oxylipin formation exhibits unique organ- and cell type-specific signatures (Schädel et al. 2021; Schadel et al. 2023), underlining the necessity to consider age as variable when investigating inflammation.

Furthermore, lifestyle crucially determines biological health and the severity of ageing (Larsson, Kaluza, and Wolk 2017; Wang, Chen, et al. 2023). Social isolation and loneliness are directly associated with poor health outcomes, increased inflammation, and cognitive decline in ageing (Leigh-Hunt et al. 2017; Cardona and Andres 2023). A meta-analysis of 148 studies on social relationships and mortality risks has found that individuals with stronger social relationships exhibit a higher likelihood of survival (Holt-Lunstad, Smith, and Layton 2010). Moreover, genome-wide analyses revealed that people experiencing a high degree of subjective social isolation show increased leukocyte transcriptional activity, especially in pro-inflammatory pathways, directly linking social isolation with inflammation (Cole et al. 2007). Maintaining social connections in old age has proven to be particularly beneficial, as the negative impact of social isolation is estimated to be higher than that of diabetes (Yang et al. 2016). Mouse studies on social isolation show elevated corticosterone levels, which acts as the main active endogenous glucocorticoid in mice, indicating a physiological stress response, along with increased inflammation and impaired cognitive and immune function (Berry et al. 2012; Ederer et al. 2022; Muta et al. 2023). Conversely, a positive social environment is associated with better immunity and increased lifespan (Garrido et al. 2022), underscoring the critical role of social connections in both ageing and inflammatory processes. But how social isolation affects oxylipin networks and which roles oxylipins play in physiological consequences of social isolation is unknown.

Metabolic interventions such as voluntary exercise (e.g., for humans taking a walk, for mice voluntary wheel running) are well established as a tool that may prolong the life and health span of an individual (Lopez-Otin et al. 2016) and subsequently may mitigate the negative impact of other lifestyle factors like isolation. While regular exercise yields a multitude of positive outcomes, the exact molecular drivers behind these benefits are not yet fully understood. Exercise not only decreases the secretion of pro-inflammatory cytokines but also shifts the oxylipin pathways toward omega-3 substrates, leading to elevated levels of SPM and their precursors (Markworth, Maddipati, and Cameron-Smith 2016; Pena Calderin et al. 2025). Additionally, it was found that exercise reduces perceived loneliness (Pels and Kleinert 2016). Although previous studies have examined inflammageing, age-related social isolation, and late-life exercise interventions individually, the combined effects and interactions of these factors remain unexplored. Here, we elucidated the implications of age-associated social isolation and related stress on the inflammatory microenvironment in aged mice, focusing on oxylipins, as well as cognitive function, and whether accompanied exercise (i.e., voluntary wheel running) can mitigate the detrimental effects of social isolation. We accounted for differential plasticity in various organs by screening seven different organ/tissue types, as well as serum (systemic circulation), creating a fundamental framework linking inflammation, ageing, and age-associated isolation stress.

## Results

The inflammatory status of 7 organs (brain, heart, fat, liver, lung, muscle, spleen) and serum of adult (5 months) and aged (20 months) C57BL6/J/UKJ mice was assessed by analysis of inflammation-related cytokine and oxylipin profiles. To investigate the effect of social isolation stress at a later stage in life, we recurrently separated mice at an age of 18 months into single cages for three individual nights per week over a period of 8 weeks. Moreover, we addressed exercise as an established, anti-inflammatory metabolic intervention by providing a running wheel to a subgroup of isolated aged mice. In this context, aged mice additionally underwent a Barnes maze test to assess intervention-related changes in cognition. The detailed experimental setup is depicted in Figure 1.

**Figure 1:**
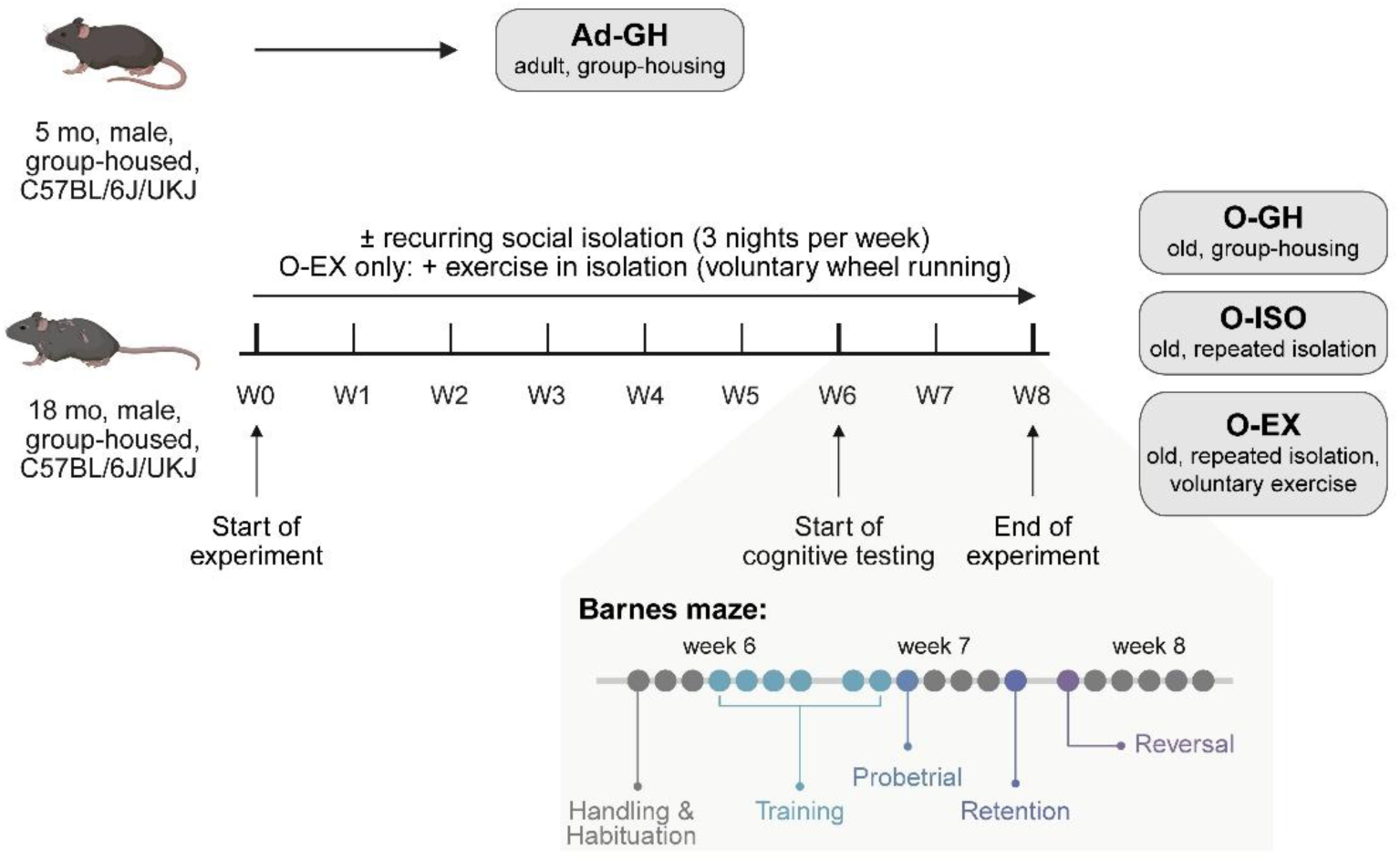
Schematic representation of the recurring social isolation and physical exercise intervention. 5-month-old, adult male mice (Ad-GH, *n* = 5) were kept in group housing until sacrificed. For the O-GH cohort, aged mice were maintained in group housing throughout their lives and underwent cognitive testing in the last three weeks before sacrifice. To investigate the effect of recurring social isolation in late-life, 18-month-old mice were repeatedly isolated into single cages for 3 individual nights per week, and resocialised with their corresponding littermates throughout a total of 8 subsequent weeks. The single cages were equipped with (= O-EX) or without (= O-ISO) running wheels. Aged mice underwent cognitive testing (*n* = 17-18) during the last weeks before the end of the experiment. Organs and blood for serum investigation (*n* = 5-6) were collected immediately after sacrifice.

### Organ-specific inflammatory cytokines are differentially affected by ageing but significantly induced by recurring social isolation

Ageing significantly increased (*p* = 0.018) the body weight of the mice. <u>O</u>ld mice in <u>g</u>roup-<u>h</u>ousing (O-GH) had a mean weight of 35.2 ± 0.6 g while the mean weight of <u>ad</u>ult mice in <u>g</u>roup-<u>h</u>ousing (Ad-GH) was 32.1 ± 0.9 g. Social isolation stress in <u>o</u>ld mice due to repeated <u>iso</u>lation into single cages (O-ISO) did not significantly alter the final body weight (35.9 ± 1.7 g, *p* = 0.692). However, the body weight of these isolated aged mice (O-ISO) showed a higher degree of variation versus old mice in group-housing (O-GH) (Fig. 2a).

**Figure 2:**
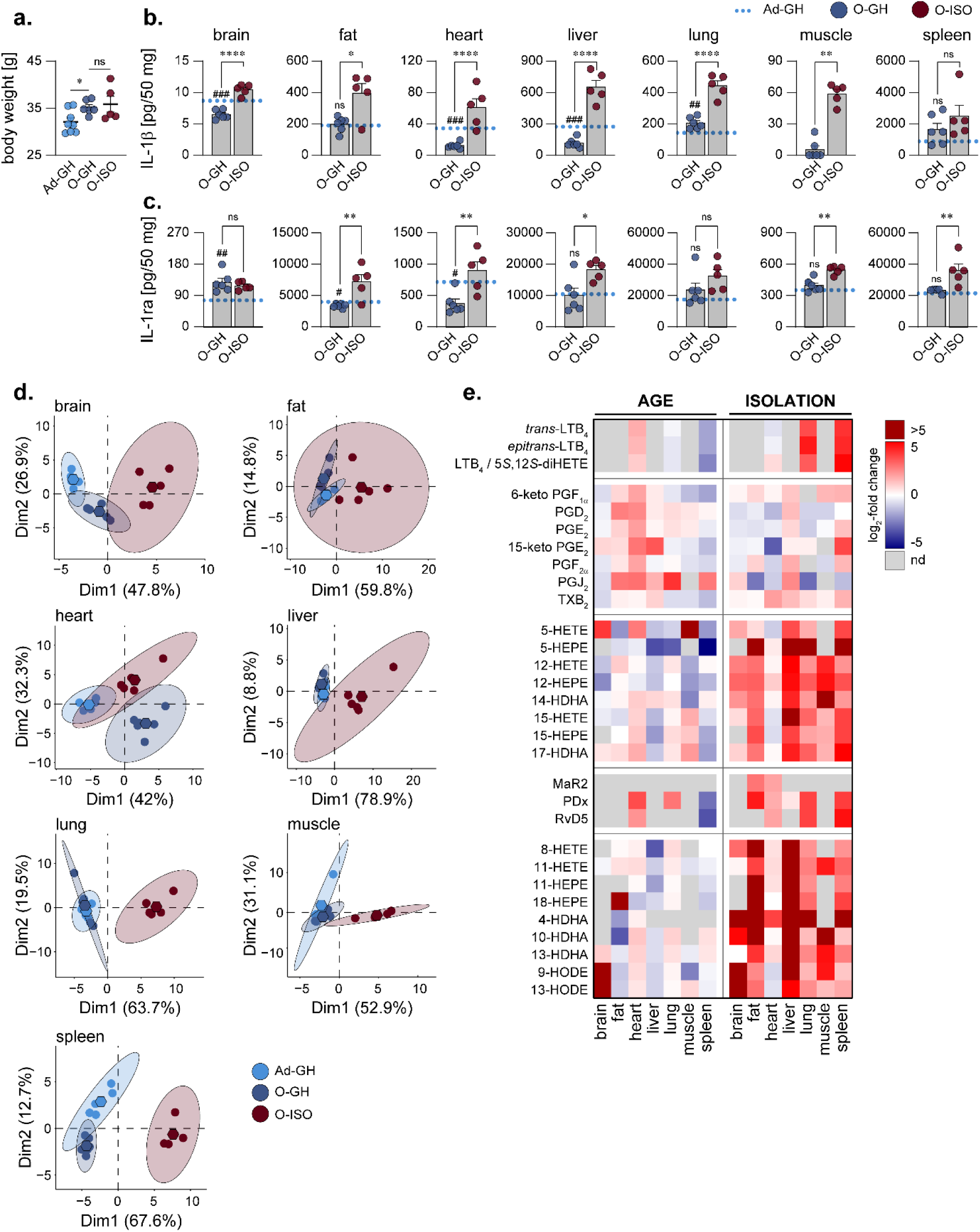
Ageing leads to organ-specific inflammatory phenotypes that are aggravated by social isolation stress. **(a)** Body weight of adult (5 months) and aged (20 months) mice after being kept in group-housing (Ad-GH and O-GH, respectively) or repeatedly isolated (O-ISO). **(b,c)** Concentration of the cytokines IL-1β and IL-1ra in pg per 50 mg organ. **(d)** Principal component analysis of organ-specific oxylipin profiles. Hexagons indicate the mean PCA score of all screened replicates within respective experimental groups. **(e)** Log_2_-fold changes for the organ-specific concentrations of individual oxylipins for the comparison of aged (O-GH) and adult (Ad-GH) mice kept in group-housing (left panel), and the comparison of aged, repeatedly isolated (O-ISO) and aged, group-housed (O-GH) mice (right panel). Fold-changes that could not be calculated due to missing values (e.g., below limit of detection = nd) are depicted in grey. **Statistics**: Data are shown as (**a**-**c**) mean ± SEM. The number of biological replicates is *n* = 5 for Ad-GH and O-ISO, *n* = 6 for O-GH. Unpaired, two-tailed Student’s *t*-tests with or without Welch-correction were performed for indicated comparisons. #, comparison of O-GH versus Ad-GH.

To address the inflammatory status of the investigated organs, we examined the protein level of the pro-inflammatory cytokine interleukin (IL)-1β, and its counteractor IL-1 receptor antagonist (IL-1ra). We found that ageing did not markedly change IL-1β levels in fat, muscle, and spleen (Fig. 2b). However, IL-1β levels were significantly decreased due to ageing in brain (Ad-GH: 8.7 ± 0.2, O-GH: 6.5 ± 0.3 pg/50 mg), heart (Ad-GH: 34.5 ± 5.3, O-GH: 12.3 ± 1.5 pg/50 mg), and liver (Ad-GH: 273.7 ± 31.7, O-GH: 117.5 ± 17.6 pg/50 mg). Only in the lung, IL-1β was significantly elevated (Ad-GH: 143.5 ± 8.1, O-GH: 205.9 ± 14.6 pg/50 mg) in the aged compared to the adult mice (Fig. 2b). When the aged mice underwent social isolation (O-ISO), however, IL-1β significantly increased in all studied organs except in spleen with only minor elevation versus the O-GH mice (Fig. 2b, 1.8-fold, *p = 0.234*). Overall, this translates to a distinct, organ-specific ageing-related impact on IL-1β secretion while stress due to isolation seems to consistently promote inflammation in most organs. A similar pattern was observed for the levels of IL-1ra, being significantly increased following isolation of aged mice in fat, heart, liver, muscle, and spleen (Fig. 2c).

### Social isolation potently induces lipoxygenase-derived oxylipin formation

Since oxylipins are crucial regulators of all stages of inflammation (Funk 2001; Levy et al. 2001; Serhan 2014) we assessed the oxylipin signature profiles of the different organs using a targeted lipidomic approach based on UPLC-MS/MS. To get an overview of age- and isolation-related changes in the complex oxylipin profiles, we performed an unbiased principal component analysis. This analysis revealed distinct age-related alterations in the oxylipin profiles for brain, fat, heart, and spleen between both group-housed adult (Ad-GH) and aged mice (O-GH) (Fig. 2d). An even more pronounced separation of the clusters for all organs was observed following isolation of aged mice (O-ISO) and only in the heart, age had a stronger effect on oxylipin profiles than isolation (Fig. 2d).

Looking at the changes of the individual screened oxylipins due to ageing, we found distinct organ-specific ageing signatures with liver, muscle, and spleen exhibiting an overall decrease in oxylipins, whereas brain, fat, and lung showed divergent oxylipin profiles with age (Fig. 2e, left). In the heart, ageing led to increased amounts of oxylipins, particularly pronounced for PGs (Fig. 2e, left). Social stress in aged mice due to isolation, however, drives a remarkable increase of a broad spectrum of oxylipins, especially derived from 5-, 12-, and 15-LOX across various organ systems (Fig. 2e, right). Interestingly, in lung and spleen, the inflammation-resolving SPMs RvD5 and PDX produced by12/15-LOX were elevated along with pro-inflammatory LTs derived from 5-LOX. In the brain and heart, the oxylipin levels were mostly unaffected by isolation of the aged mice (Fig. 2e, right).

In order to further characterise the age- and isolation-related effects on the oxylipin formation, we classified the oxylipins according to their rate-limiting biosynthetic enzymes, i.e., COX, 5-LOX, 12-LOX, and 15-LOX. In detail, for COX products, age-related increases were most pronounced in the heart (3.71-fold) with minor changes in other organs. Isolation did not affect COX products in most organs (fat, heart, lung, and muscle), except for brain (0.71-fold), liver (2.79-fold) and spleen (1.41-fold; Fig. 3a). Taken together, the effect size and its statistical significance indicate that age and isolation only have a moderate impact on COX-derived oxylipins in most organs.

**Figure 3:**
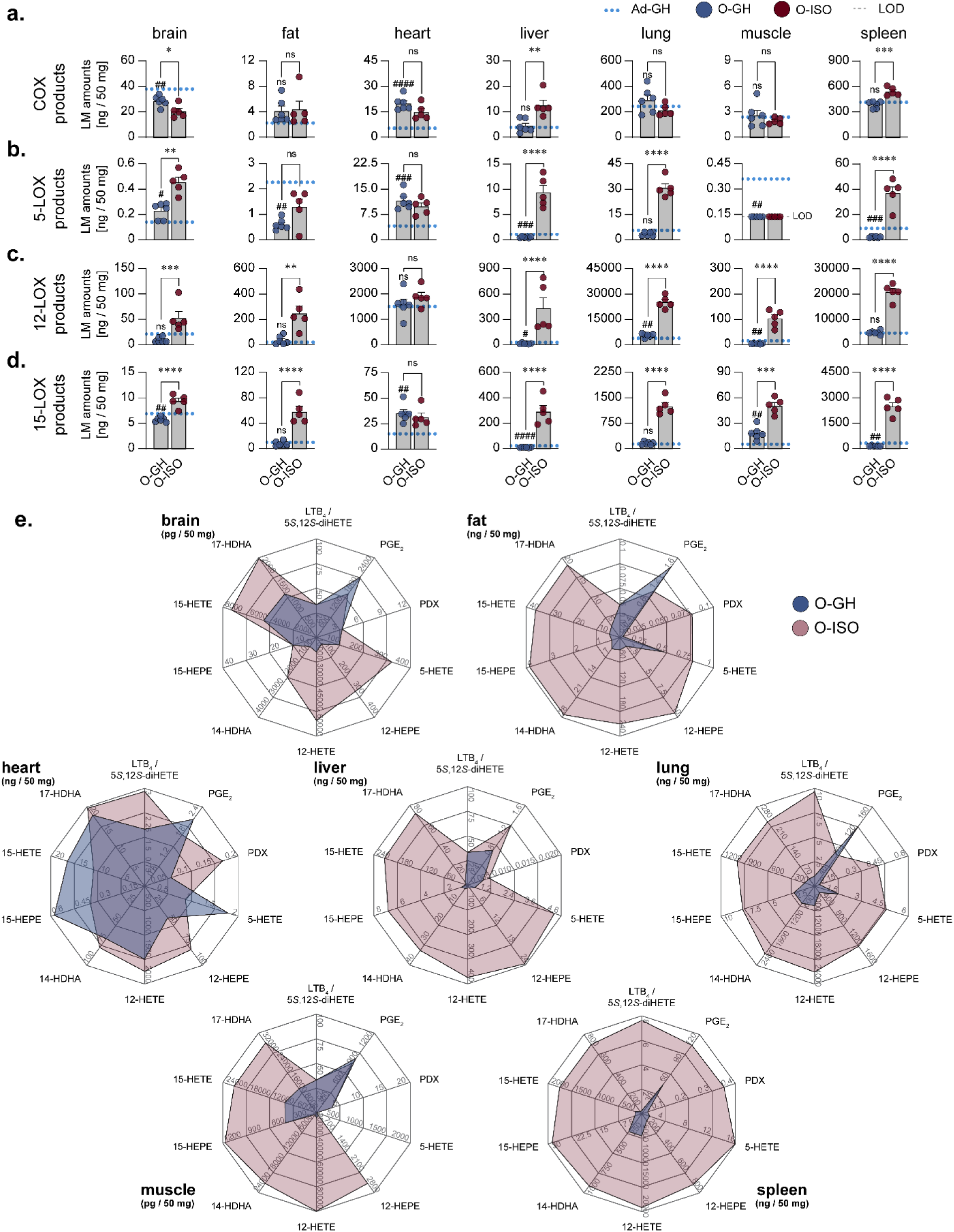
Social isolation stress leads to induction of LOX pathways. Total grouped amounts of oxylipins that are products of the COX **(a)** or LOX **(b-d)** pathways. Metabolites were grouped as follows: *COX* – PGD_1_, PGE_1_, PGF_1α_, 6-keto PGF_1α_, PGD_2_, PGE_2_, 15-keto PGE_2_, PGF_2α_, PGF_2ß_, PGJ_2_, PGD_3_/PGE_3_, PGF_3α_, TXB_2_; *5-LOX* – RvE1, RvE2, *trans*-LTB_4_, *epitrans*-LTB_4_, LTB_4_/5*S*,12*S*-diHETE, 5*S*,6*R*-diHETE, 20-OH LTB_4_, LTB_5_, 5-HETE, 5-HEPE, 7-HDHA; *12-LOX* – MaR1, MaR2, 12-HETE, 12-HEPE, 14-HDHA; *15-LOX* – PDx, PD1, RvD1, RvD2, RvD3, RvD4, RvD5, RvE4, LXA_4_, LXB_4_, LXA_5_, 5*S*,15*S*-diHETE, 15-HETE, 15-HEPE, 17-HDHA. Values are given as ng per 50 mg organ. LOD of the oxylipins is indicated, if applicable. **(e)** Radar charts of individual oxylipins in aged, group-housed mice (O-GH, blue) and aged mice undergoing repeated isolation (O-ISO, red). **Statistics**: Data are shown as (a-d) mean ± SEM. The number of biological replicates is *n* = 5 for Ad-GH and O-ISO, *n* = 6 for O-GH. Unpaired, two-tailed Student’s *t*-tests with or without Welch-correction were performed for indicated comparisons. #, comparison of O-GH versus Ad-GH.

5-LOX products were much more affected by age and isolation. Thus, 5-LOX product levels strongly decreased in fat, liver, muscle, and spleen of aged mice compared to adult controls (Fig. 3b). Conversely, ageing markedly increase 5-LOX products in the heart, which is surprisingly not further aggravated by isolation (Fig. 3b). For all other organs – except muscle where 5-LOX products were not detectable – isolation led to a marked increase of 5-LOX-derived oxylipins (Fig. 3b, brain: 2.0-fold, fat: 2.1-fold, liver: 18.2-fold, lung: 8.8-fold, spleen: 17.1-fold). The same trend was observed for oxylipins produced by 12- and 15-LOX (Fig. 3c,d). Thus, ageing caused only moderate changes in the levels of 12-LOX and 15-LOX products, while social isolation caused a dramatic increase for both 12-LOX (brain: 5.7-fold, fat: 7.1-fold, liver: 30.1-fold, lung: 4.5-fold, muscle: 22.2-fold, spleen: 4.3-fold) and 15-LOX products (brain: 1.6-fold, fat: 7.5-fold, liver: 33.4-fold, lung: 7.9-fold, muscle: 2.8-fold, spleen: 15.6-fold) in all organs except for the heart (no elevation). In more detail, the radar charts showing individual representative oxylipins for each enzyme class (COX: PGE_2_; 5-LOX: LTB_4_ and 5-HETE; 12-LOX: 12-HEPE, 12-HETE and 14-HDHA; 15-LOX: PDX, 15-HEPE, 15-HETE and 17-HDHA) confirm the massive elevation of LOX-derived oxylipins in all organs caused by social stress due to isolation, except for heart, with minor or even opposite impact on COX products (e.g., PGE_2_). Together, social isolation stress consistently upregulates LOX-derived oxylipins in organs of aged mice while only the heart and, to a lesser extent, the brain exhibit a lower susceptibility to social stress-related oxylipin alterations (Fig. 3e). These increases in LOX-derived oxylipins were evident for conversion of all PUFAs as substrates, i.e. AA, EPA and DHA.

### Late-life exercise cannot rescue isolation effects on inflammatory state

Since voluntary exercise yields a multitude of positive outcomes prolonging the life and health span (Lopez-Otin et al. 2016) with impact on cytokines and oxylipins (Markworth, Maddipati, and Cameron-Smith 2016; Pena Calderin et al. 2025), we further investigated whether exercise can mitigate the effects on the inflammatory mediators induced by social isolation stress. We employed voluntary wheel running as a late-life exercise intervention in socially isolated aged mice (runners: O-EX, non-runners: O-ISO). After the intervention period, the group of O-ISO mice (non-runners) showed a mean weight of 35.9 ± 1.7 g, whereas the group of O-EX mice (runners) showed a slight decrease in body weight (33.9 ± 1.1 g), yet this difference was not significant (Fig. 4a, *p* = 0.357). However, the weight loss seems to be connected to the individual running performance of the mice, as the mouse with the lowest running performance across the 10-week-long experimental period showed the highest weight and vice versa (Fig. 4b).

**Figure 4:**
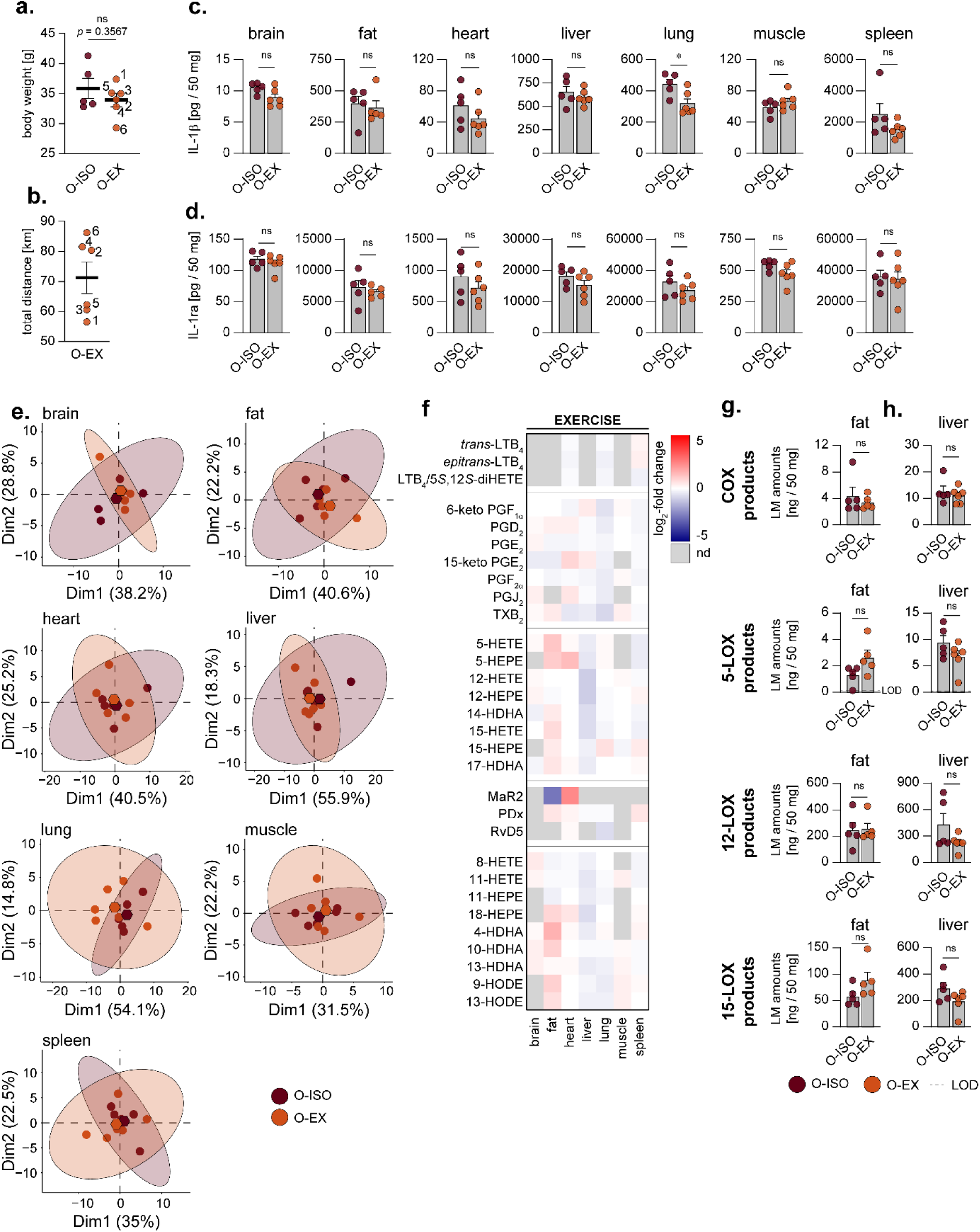
Exercise cannot ameliorate increased oxylipin secretion in aged mice affected by social isolation stress. **(a)** Body weight of aged (20 month) mice after being repeatedly isolated into single cages with (O-EX) or without running wheels (O-ISO) for 3 separate nights per week over a period of 8 weeks. **(b)** Total running performance of O-EX mice is summarised in km over the experimental period of eight weeks. **(a,b)** Numbers indicate the individual animals in the O-EX cohort. **(c,d)** Concentration of IL-1β and IL-1ra in pg per 50 mg organ from O-ISO and O-EX mice. **(e)** Principal component analysis of organ-specific oxylipin profiles from O-ISO and O-EX mice. Hexagons indicate the mean PCA score of all screened replicates within respective experimental groups. **(f)** Log_2_-fold changes for the organ-specific levels of individual oxylipins comparing the levels of O-EX versus O-ISO mice. Fold-changes that could not be calculated due to missing values (e.g., below limit of detection = nd) are depicted in grey. **(g,h)** Total amounts of grouped oxylipin species derived from the COX or LOX pathways in **(g)** fat and **(h)** liver. Oxylipins were grouped as indicated in Fig. 3. Values are given as ng per 50 mg organ. Limit of detection (LOD) of oxylipins is indicated, if applicable. **Statistics**: Data are shown as (**a-d**, **g-h**) mean ± SEM. The number of biological replicates is *n* = 5 for O-ISO and *n* = 5-6 for O-EX. Unpaired, two-tailed Student’s *t*-tests with or without Welch-correction were performed for indicated comparisons.

We then investigated the levels of IL-1β and IL-1ra and assessed the oxylipin signature profiles of organs from aged mice during isolation that were either allowed to exercise (O-EX) or not (O-ISO). Only in the lung, exercise led to a significant decrease in IL-1β levels (0.72-fold) (Fig. 4c) and a small decrease by trend was observed in heart (0.72-fold, *p* = 0.241) and spleen (Fig. 4c, 0.62-fold, *p* = 0.151). For all other organs, the IL-1β levels were essentially unchanged. Similarly, IL-1ra levels remained essentially unaltered between O-EX and O-ISO, without significant impact of exercise in any organ (Fig. 4d). Principal component analysis of the overall oxylipin profiles showed no marked differences (no separation of clusters) for all organs comparing O-ISO versus O-EX, with both the position of mean and 95% confidence intervals being similar (Fig. 4e). Yet, when looking at individual oxylipins in detail (Fig. 4f), oxylipin levels in fat (increased) and liver (decreased) were most affected by exercise, particularly pronounced for monohydroxylated fatty acids like 5-HETE, 4-HDHA, 12-HETE, 12-HEPE etc. (Fig. 4f). Interestingly, in the lung, exercise decreased PG and RvD5 formation but did not affect other oxylipins (Fig. 4f). Analysis of oxylipin groups according to biosynthetic enzymes (i.e., COX, 5-, 12-, 15-LOX) reveals no significant alterations due to exercise in any organ (Fig. S2a-d), and only in fat and liver a tendency for changes was observed. Thus, the most notable change by trend upon exercise is the increase of 5-and 15-LOX-derived products in fat tissue (5-LOX: 2.02-fold, *p* = 0.131; 15-LOX: 1.52-fold, *p* = 0.082) while COX- and 12-LOX-derived products remain at a similar level to sedentary mice (Fig. 4g). Furthermore, exercise led to a small decrease in LOX-derived products (5-LOX: 0.72-fold, *p* = 0.214; 12-LOX: 0.52-fold, *p* = 0.140; 15-LOX: 0.67-fold, *p* = 0.188) in the liver, while the level of COX-derived products was unaltered (Fig. 4h). Together, for aged mice undergoing social isolation stress, the variable of voluntary exercise has only a minor impact on the massive alterations of oxylipins and cytokine levels caused by isolation.

### Cognitive function does not benefit from voluntary exercise in isolation

In a previous study (Ederer et al. 2022), we addressed the relation between age and cognition in male, group-housed mice by subjecting them to a Barnes maze test, revealing a cognitive decline that occurs between the ages of 15 and 24 months. It was also shown that for aged mice, constant social isolation further impairs cognitive function. To evaluate whether voluntary wheel running can mitigate cognitive decline in ageing and social stress through repeated isolation and resocialisation, we tested isolated aged mice in the Barnes maze to assess spatial learning, as well as short- and long-term memory. The timeline of the implementation of the Barnes maze test is shown in the schematic in Fig. 5a. During the training period for the Barnes maze test, both cohorts with recurring social stress (O-ISO and O-EX) showed a similar, improved performance over time. No strong exercise-mediated improvements were observed during training (Fig. 5b). The probe trial (short-term memory) and the retention test (long-term memory) did not reveal cognitive improvement following exercise (Fig. 5c,d). During the reversal test (cognitive flexibility), the O-EX mice showed a shorter primary latency and an increased cognitive score compared to O-ISO mice, suggesting a potential facilitation in task adaptation (Fig. 5e). Overall, exercise intervention in late life does not seem to improve spatial learning, short and long-term memory in aged mice experiencing social isolation stress, but O-EX showed an improved cognitive flexibility.

**Figure 5:**
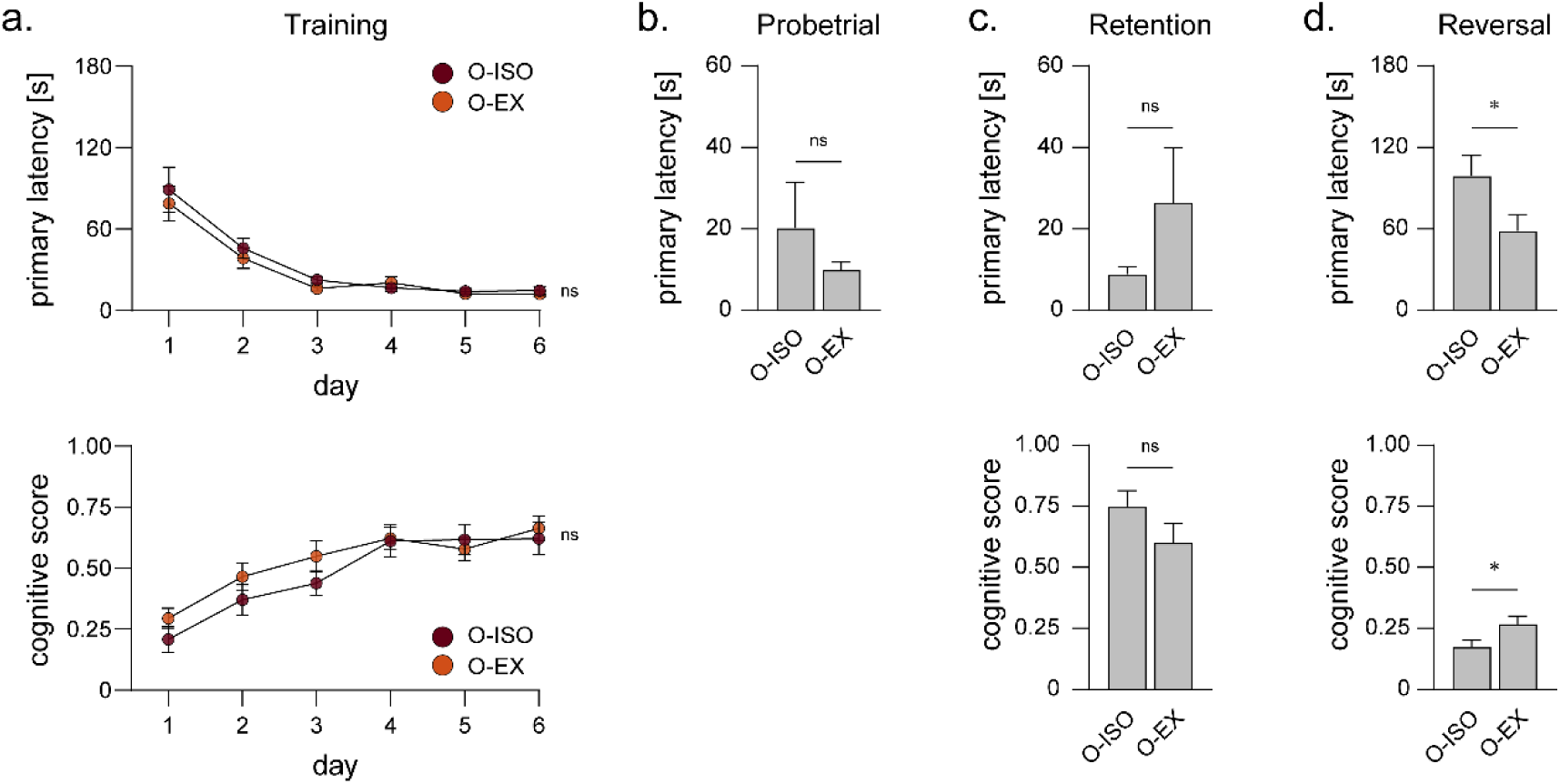
Cognitive function assessment of mice undergoing recurring social isolation with and without physical exercise. Results from the Barnes maze test, depicted as the primary latency period and as cognitive scores for **(b)** training, **(c)** probe trial, **(d)** after a retention period, and **(e)** in a reversal test. **Statistics**: Data are shown as mean ± SEM. The number of biological replicates is *n* = 18 for O-GH and *n* = 17 for O-EX. Unpaired, two-tailed Student’s *t*-tests with or without Welch-correction were performed for indicated comparisons.

### Reduced age-associated inflammatory repertoire of oxylipins and proteins in serum is maintained after isolation and exercise

To determine whether the observed organ-specific effects due to ageing and isolation as well as exercise are reflected in the systemic circulation, we characterised the inflammatory secretome (oxylipins and proteins) in the serum of the mice cohorts from above. Based on both, the principal component analysis of the oxylipin profiles and the detailed analysis of the individual oxylipins, age but not isolation or exercise is the main driver for the observed alterations (Fig. 6a,b). While in the PCA all aged cohorts (O-GH, O-ISO, O-EX) cluster tightly together, the oxylipin profile of the adult cohort (Ad-GH) shows a distinct shift (Fig. 6a). The majority of the screened oxylipins in serum are decreased due to age (O-GH/Ad-GH, Fig. 6b) with significant changes for 5- and 15-LOX-derived products (5-LOX: 0.55-fold, 15-LOX: 0.54-fold, Fig. 6c). To our surprise and in contrast to organs (see above), isolation only had a minor impact on oxylipin levels in serum, reflected by the lack of changes for COX and 12-LOX products, and only moderate increases in 5- and 15-LOX-derived products in serum from O-ISO versus O-GH mice (5-LOX: 1.63-fold, *p* = 0.014; 15-LOX: 1.44-fold, *p* = 0.174, Fig. 6c). These serum oxylipin levels of O-ISO did not markedly change due to exercise, apart from a 4-fold increase in 5-LOX-derived products (O-ISO: mean = 822.2 ± 124.0 pg/mL, O-EX: mean = 3341.3 ± 1661.3 pg/mL; *p* = 0.211) which was, however, not significant and attributed to only 2 out of the 6 biological replicates (Fig. 6c).

**Figure 6:**
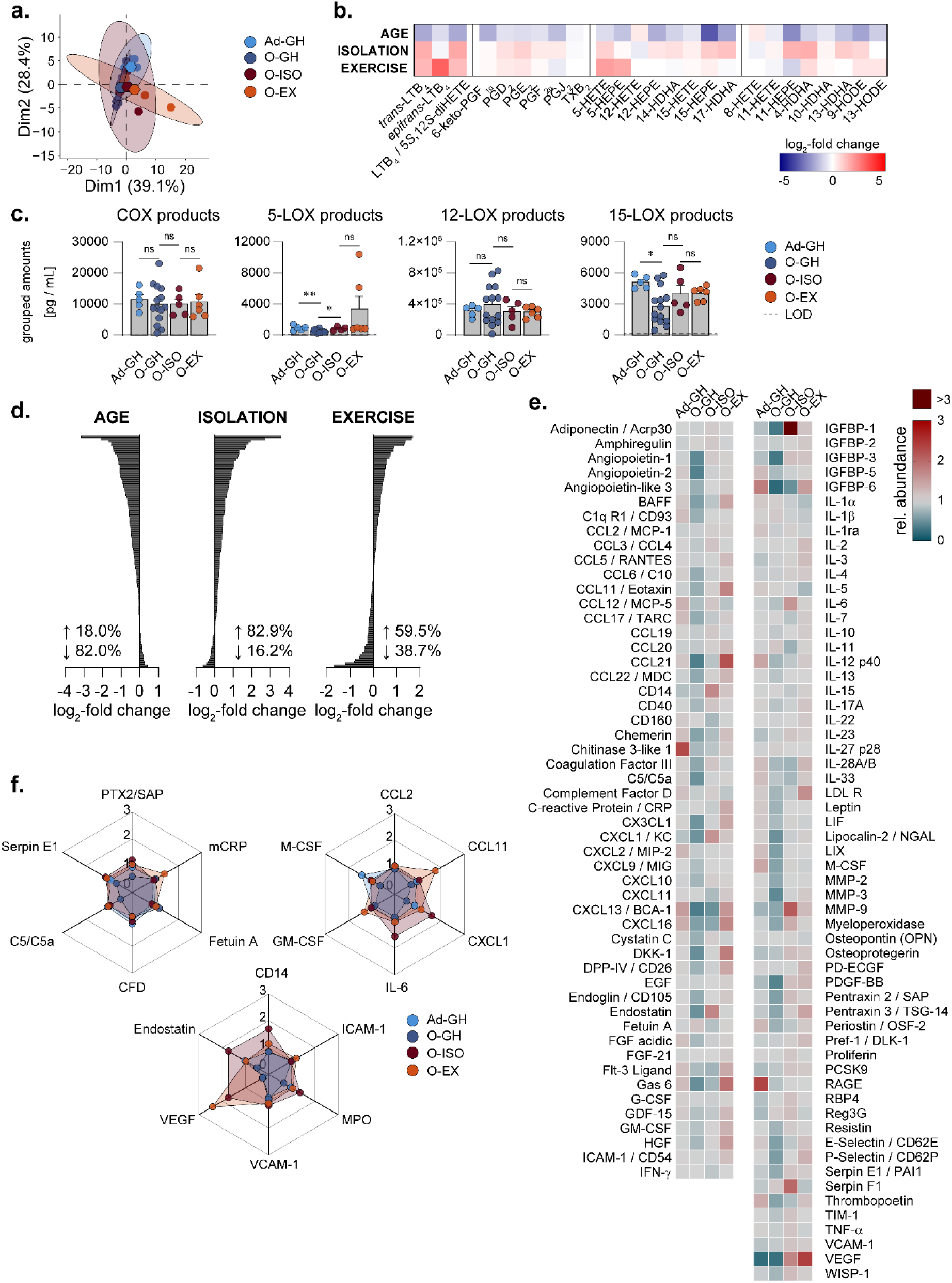
Impact of age and social isolation on inflammatory oxylipins and proteins in serum. **(a)** Principal component analysis of oxylipin profile of serum. Hexagons indicate the mean PCA score of all screened replicates within respective experimental groups. **(b)** Log_2_-fold changes for the concentrations of individual lipid mediators, detected in serum, comparing the levels of (AGE) O-GH versus Ad-GH, (ISOLATION) O-ISO versus O-GH, and (EXERCISE) O-EX versus O-ISO mice. **(c)** Total amounts of grouped LM species that are products of the COX or LOX pathways in serum. Metabolites were grouped as indicated for Fig. 3. Values are given as pg per mL serum. Limit of detection (LOD) of the metabolites is indicated, if applicable. **(d)** Log_2_-fold changes of 111 screened, circulating proteins in pooled serum samples and percentages of regulated proteins for the depicted comparisons. **(e)** Heatmap showing the relative abundance of the investigated proteins in all experimental cohorts. **(f)** Radar charts displaying the relative abundance of selected (top left) acute-phase proteins, (top right) classical cytokines and chemokines, and (bottom) various surface markers and inflammatory proteins. **Statistics**: Data are shown as mean ± SEM (**c**). The number of biological replicates is *n* = 5 for Ad-GH and O-ISO, *n* = 14 for O-GH and *n* = 6 for O-EX. (**d-f**) Samples were pooled from the serum of 4-5 biological replicates for all screened cohorts. Unpaired, two-tailed Student’s *t*-tests with or without Welch-correction were performed for indicated comparisons.

We next employed an unbiased analysis of the repertoire of inflammation-related cytokines, chemokines, enzymes, and growth factors in the serum of Ad-GH, O-GH, O-ISO, and O-EX mice, using a commercially available Proteome Profiler, allowing us to screen for the relative abundance of 111 candidates. In line with the reduced amounts of most of the oxylipins (Fig. 6b), ageing led to an overall downregulation (82.0%) of the investigated proteins in serum (Fig. 6d). Isolation, however, led to an increased expression of 82.9% of the proteins (Fig. 6d). Despite the seemingly contradictory dynamics, the proportions of proteins being down-regulated with age and up-regulated due to isolation are not congruent (Fig. S3). The exercise intervention (O-EX) led to less pronounced and more balanced alterations for aged mice under isolation (O-ISO) with 59.5% up-and 38.7% downregulated proteins (Fig. 6d). Acute phase proteins (APPs, e.g., complement factors and pentraxins) and chemokines (particularly of the CCL and CXCL classes) were impaired due to age versus adult mice (Fig. 6e,f). Such overarching class effects could not be observed for the cohorts O-ISO and O-EX under isolation. Interestingly, the pro-inflammatory cytokine IL-6, the chemokine CXCL1, and the LPS receptor cluster of differentiation (CD)14 were increased in O-ISO versus O-GH mice (Fig. 6e,f). Exercise of aged mice under isolation, however, induced higher levels of murine C-reactive protein (CRP), CCL11, CCL21, and vascular endothelial growth factor (VEGF) versus O-ISO mice (Fig. 6e,f). These alterations in the abundance of inflammation-related serum markers indicate a reduced immune competence with age, and distinct pro-inflammatory responses due to isolation and exercise.

## Discussion

Social isolation, a phenomenon commonly occurring in late life, is extensively described as a risk factor for increased morbidity and mortality and is associated with higher levels of inflammation and cognitive decline (Hodgson et al. 2020; Wang, Gao, et al. 2023). Considering inflammation as a shared feature of ageing (Ferrucci and Fabbri 2018), isolation (Yang et al. 2013; Smith et al. 2020), and an inactive lifestyle (Fenton et al. 2018), we examined the inflammatory cytokine and oxylipin secretome in mice undergoing recurring social isolation and resocialisation, and whether voluntary physical exercise can ameliorate the inflammatory stress responses due to isolation in single housing. Unexpectedly, we found dramatic elevation of inflammation-related LOX-derived oxylipins due to social isolation in an organ-specific manner, especially affecting liver, lung, and spleen. Surprisingly, physical exercise showed a neglectable impact on the organ-intrinsic oxylipin secretome of isolated mice and was hardly evident in systemic circulation.

Previous studies have frequently investigated and described social isolation as a stressor, resulting in a variety of adverse health outcomes (Grant, Hamer, and Steptoe 2009; Mumtaz et al. 2018). The magnitude of these events, however, varies greatly depending on the type and duration of social isolation (Shoji and Mizoguchi 2011; Hammig 2019). Studies on permanent or short-to long-term social isolation reported mixed and sometimes contradictory results. Thus, a four-week social isolation period of both male and female adult mice (3 months) affected female animals more than males, but overall, the isolation only had minimal effects on serum levels of the stress marker corticosterone, behaviour, and body weight (Smolensky et al. 2024). Conversely, shorter periods of single housing of mice, such as 4 - 5 days only, tend to produce more pronounced detrimental effects, including anxiety behaviour, elevated urinary stress hormones, and weight loss (Muta et al. 2023). These findings suggest that, while isolation quickly triggers a strong stress response, mice may adapt within a few weeks, resulting in lower or unchanged stress markers over prolonged isolation. Nevertheless, despite seemingly modest stress responses, long-term isolation of degus over a period of 20-24 months still promotes a pro-inflammatory state (Oliva et al. 2023). Research on the effects of long-term isolation in human cohorts is limited. One study reported elevated cortisol levels and heightened immune activation in men after an isolation period of approximately 1.5 years (Yi et al. 2014). However, to allow comparability between studies on isolation, further adjustments to the experimental conditions are needed. In our study, aged mice underwent a social stress protocol of repeated isolation in single housing for three individual nights per week and subsequent resocialisation in group housing over a total period of 8 weeks. Our results confirm that such repeated social isolation acts as a potent and impactful stressor, elevating the pro-inflammatory cytokine IL-1β. We showed before, that oxylipins are affected by ageing in an organ-specific manner (Schädel et al. 2021), which we also confirm in this study. Additionally, we here uncover a dramatic upregulation of LOX-derived oxylipin production in several organs due to isolation, while COX products that are produced from the same PUFA substrates (Funk 2001) were hardly altered. Increased IL-1β has previously been linked to an isolation-associated inflammatory phenotype (Magalhaes et al. 2024), but our findings demonstrate, for the first time the tremendous effect of social isolation stress on oxylipins as responsible mediators of the inflammatory response in several organs. Notably, these age- and isolation-related effects did not occur systemically, reflected by the overall moderate changes of oxylipin levels in serum of mice.

The dramatic changes in oxylipin formation due to isolation may be attributed to a dysregulation of the hypothalamic-pituitary-adrenal (HPA) axis in response to social isolation as a psychosocial stressor, resulting in elevated glucocorticoid (GC) signalling. It is known that oxylipins, particularly eicosanoids such as PGs, can be affected by stress-related GC (Umamaheswaran et al. 2018), and that GC may influence the expression of COX-1 and −2 (Goppelt-Struebe, Wolter, and Resch 1989; Ehrchen et al. 2007). LOX expression has likewise been shown to be regulated through exogenous GC, which can decrease the pro-inflammatory immune response along with elevated 15-LOX-2 expression and elevated SPM levels (Riddick et al. 1997; Rao et al. 2023). Although the murine paralog of 15-LOX-2, also called m8-LOX, is known to oxygenate free substrates at the 8-position primarily, it has been demonstrated to oxygenate the 15-position in the case of phospholipid-esterified fatty acid substrates (Bender et al. 2016). This could explain the observed increase in 15-LOX-derived products. Our present results documenting elevated LOX products in several organs and tissues of aged mice due to social isolation stress, predominantly in immunological organs like spleen, liver, and lung, are in good agreement with the above-mentioned impact of GC on oxylipin biosynthesis. Whether or not isolation of aged mice elevates LOX expression and if GC are involved in this process remains to be explored in more detail.

Aberrations in inflammatory signalling caused by age and isolation may raise the risk of dysregulated homeostasis, leading to chronic basal inflammation. Cognitive decline may also result from age- and stress-related imbalances in oxylipins, as PG signalling and 5-LOX expression have been associated with reduced cognitive function and depression-like symptoms (Luo et al. 2016; Minhas et al. 2021; Mrowetz et al. 2023). Furthermore, chronic social stress has been found to significantly elevate levels of 12-LOX-derived metabolites 12-HETE and 12-HEPE in the brain (Akiyama et al. 2022). In our study, isolation of aged mice elevated both, 5-LOX products with mainly pro-inflammatory features (Haeggstrom 2018) but also 12/15-LOX-derived SPM like RvD5, PDX, and MaR2 and their precursors 17-HDHA and 14-HDHA, which act as immunoresolvents that terminate and resolve inflammation (Serhan 2014). Therefore, these alterations in oxylipin profiles may be connected to a stress-induced activation of the innate immune system, triggering the mobilisation of neutrophils, monocytes, and monocyte-derived macrophages, as primary producers of oxylipins (McKim et al. 2018).

Lifestyle interventions such as diet or exercise are popular and well-characterised means of mitigating or circumventing negative age-related health outcomes (Ngandu et al. 2015; Fitzgerald et al. 2021). Exercise has been shown to exhibit anti-inflammatory features, which are, however, highly dependent on type, duration, onset, and intensity (Gleeson et al. 2011). For example, Sun et al. found that exercise indeed reduces inflammageing features like IL-1β secretion during long-term exercise (12-month voluntary exercise) in young and aged mice exposed to infectious injury (Sun et al. 2023). Thus, mice that underwent exercise were more protected from LPS challenge than sedentary mice. However, this effect was less pronounced in mice that started the exercise regimen at an older age (Sun et al. 2023), and is in line with results indicating that adaptive and acute responses to physical activity decline with advancing age (Durham et al. 2010). This may also explain why our results did not confirm previous findings showing that exercise stimulates both pro-as well as anti-inflammatory pro-resolving lipid mediators (Calderin et al. 2022; Malan et al. 2024; Pena Calderin et al. 2025). In contrast to these studies, our exercise protocol started in late life (18 mo), in addition to the stress factor of repeated social isolation and resocialisation. We suppose that the detrimental effects of social isolation override the benefits of physical exercise, and thus, no ameliorating effect of exercise on pro-inflammatory signatures could be observed.

While our protocol of repeated isolation and resocialisation may better mimic isolation in the elderly than current long-term isolation models, several limitations should be recognised. Past studies have shown that the effects of social isolation occur in a sex-specific manner, with female mice generally being more affected than males (Smolensky et al. 2024). To understand the origin and implications of the proposed effects, future studies in female mice, ideally in comparison to male counterparts, will be essential. Additionally, inclusion of an infection model, e.g., bacterial pneumonia caused by *S. pneumoniae*, a common infection in the elderly (Gutierrez et al. 2006; Ruiz et al. 2014), would be helpful to assess not only the basal inflammatory microenvironment but also to evaluate immune competence and whether repeated social isolation stress may significantly impair immunological resilience and resolution processes.

In future studies, the molecular signalling circuits that connect social isolation and inflammageing need to be further explored. It is also important to recognise that the idea of gradual ageing is increasingly challenged, with findings indicating that nonlinear ageing may be more common and that abrupt changes might amplify ageing processes (Shen et al. 2024; Olecka et al. 2024; Olecka, Morrison, and Hoffmann 2025). As such, social isolation stress in elderly may push individuals past a critical threshold, triggering escalation of inflammation and accelerating ageing.

Our findings demonstrate that repeated cycles of social isolation and resocialisation may act as stressors, increasing inflammation-related cytokine secretion and dramatically enhancing the formation of LOX-derived oxylipins in defined organs, impacting inflammation and tissue homeostasis. Notably and to the best of our knowledge, this is the first report revealing organ-specific oxylipin modulation upon social isolation, offering a novel framework for future interdisciplinary research at the intersection of inflammation, ageing, and social isolation. Under our experimental conditions, voluntary physical exercise did not counteract the consequences of social isolation stress on oxylipin production. Conclusively, the dramatic elevation of LOX-derived oxylipins highlights the impact of social isolation stress, a common phenomenon in the elderly, on inflammatory mediators within several specific organs and propose oxylipins as determinants of stress-related inflammageing.

## Methods

### Animals and sample isolation

Male C57BL/6J/UKJ mice were bred and housed at the Central Experimental Animal Facility at Jena University Hospital, Jena, Germany. Mice were kept on a 14/10 h light/dark cycle and at 22 ± 2 °C with a relative humidity of 55 ± 10%. The mice had unrestricted access to water and food (ssniff mouse V1534-300, ssniff Spezialdiäten GmbH, Soest, Germany). Four experimental cohorts were established: (1) Ad-GH (*n* = 5), adult mice at 5 months of age that were kept in group-housing throughout their lives; (2) O-GH (*n* = 6), 20-months-old mice that were kept in group-housing throughout their lives; (3) O-ISO (*n* = 5 for oxylipin analysis, *n* = 18 for cognitive testing), aged mice that were kept in group-housing until 18 months of age, then over a period of 8 weeks, repeatedly separated for three individual nights per week (Sunday, Tuesday, Thursday) into single cages before being returned to their littermates; and (4) O-EX (*n* = 6 for oxylipin analysis, *n* = 17 for cognitive testing), aged mice treated identically to O-ISO mice but provided with running wheel during the isolation nights, allowing for voluntary wheel running.

After the experimental period, mice were euthanised by cervical dislocation. Whole blood was collected immediately after death. To extract serum, blood was kept at room temperature for around 60 min to ensure clotting. Samples were subsequently centrifuged for 30 min at 2,000 x*g*. Serum was transferred into fresh reagent tubes and stored at −80 °C. Seven organs, namely brain, fat (visceral WAT), heart, liver, lung, muscle (quadriceps and gastrocnemius), and spleen were harvested, washed with PBS (pH 7.4, SERVA, Heidelberg, Germany; 47302.03), and stored at −80 °C.

### Cognitive testing with Barnes maze

The Barnes maze is employed to assess spatial learning, short- and long-term memory, memory retrieval, and cognitive flexibility by training animals to associate distal cues with a fixed escape box (Ederer et al. 2022). Mice are placed on a brightly lit circular platform with 20 holes, one of which provides access to an escape box. The protocol comprises one day of habituation, followed by 6 training days (3 trials/day with an inter-trial interval of 60 min) to assess learning. The following day, a probe trial, for which all holes are closed, evaluates short-term memory. After three days without further training, a retention test assesses long-term memory retention with the original hole with the escape box opened again. Lastly, one day later the position of the escape box is placed 180° opposite to the original position. This reversal test measures cognitive flexibility.

### Organ homogenisation

For fat, liver, lung, muscle and spleen, organ homogenates were prepared by weighing 20-40 mg of organ tissue in tubes containing around 100 mg of lysing matrix D per 10 mg organ (M.P. Biomedicals, Irvine, CA, USA; 116540434). The lysis buffer was added at a ratio of 20 µL per milligram organ (1% (*v*/*v*) NP-40 (AppliChem, Darmstadt, Germany; A1694), 1 mM sodium orthovanadate (AppliChem; A2196), 10 mM sodium fluoride (AppliChem; A3904), 5 mM sodium pyrophosphate (Sigma Aldrich, St. Louis, MO, USA; S8282), 25 mM β-glycerophosphate (Sigma Aldrich; G9422), 5 mM EDTA (AppliChem; A2937), 25 µM leupeptin (Sigma Aldrich; L2884), 3 mM soybean trypsin inhibitor (Sigma Aldrich; T9128) and 1 mM phenylmethanesulfonyl fluoride (Sigma Aldrich; P7626)). For the homogenisation of brain and heart, one hemisphere and the whole heart, respectively, were weighed into tubes containing lysing matrix D, and 5 µL of lysis buffer per milligram organ was added. All organs were homogenised using a FastPrep-24™ 5G bead beating homogeniser using established protocols (M.P. Biomedicals).

### Cytokine quantification via ELISA

For cytokine quantification, homogenates were centrifuged at 21,100 x*g*, 4 °C for 10 min. The supernatant was transferred into fresh reagent tubes and stored at −20 °C until further analysis. The final concentration of the homogenates was 50 mg/mL for fat, liver, lung, muscle, spleen, and 200 mg/mL for brain and heart.

Quantification of cytokines in organ homogenates was performed using commercially available ELISA kits for murine IL-1β (DY401-05) and IL-1ra (DY406-15), manufactured by R&D Systems (Minneapolis, MN, USA). The assays were completed per the manufacturer’s instructions. Quantification was based standard curves generated individually for each experiment.

### Oxylipin quantification via UPLC-tandem mass spectrometry

For oxylipin quantification, immediately after homogenisation, homogenates were mixed 1:1 with ice-cold methanol (fisher chemical; 10653963). Samples were kept on ice for around 20 min, and then centrifuged at 21,100 x*g*, 4 °C for 10 min. The amount of supernatant equivalent to 20-30 mg organ was then transferred into glass vials containing 990 µL ice-cold methanol and 10 µL of deuterium-labelled internal standard [200 nM *d*_8_-5*S*-hydroxyeicosatetraenoic acid (HETE) (Cayman Chemical; 334230), *d*_4_-LTB_4_ (Cayman Chemical; 320110), *d*_5_-LXA_4_ (Cayman Chemical; 24936), *d*_5_-RvD2 (Cayman Chemical; 11184), *d*_4_-PGE_2_ (Cayman Chemical; 10007273) and 10 μM *d*_8_-AA (Cayman Chemical; 390010). Finally, lysis buffer was added to yield a total volume of 3 mL and stored overnight at −20 °C to ensure protein precipitation.

After centrifugation (1,200 x*g*, 4°C for 10 min), supernatants were transferred to clean glass vials and acidified by addition of 9 mL of MilliQ water (pH 3.5). Solid phase extraction of organ samples was performed as previously described (Werner et al. 2019). Purified samples were evaporated under continuous N_2_ flow, and the residue was resuspended in 200 µL of an equal mixture of methanol and water (VWR Chemicals, 83645320). After final centrifugation at 21,100 x*g*, 4°C for 5 min, the supernatant was used for ultra-performance liquid chromatography coupled with tandem mass spectrometry (UPLC-MS/MS) analysis. Analytes were separated on an Acquity UPLC system (Waters, Milford, MA, USA) equipped with an Acquity UPLC BEH C18 column (1.7 μm, 2.1 mm × 100 mm; Waters, Eschborn, Germany) and a pre-column of identical material with column temperature set to 50°C. The gradient was adjusted as listed in Table 1. Analytes were detected using a QTRAP 5500 mass spectrometer (ABSciex, Darmstadt, Germany) with electrospray ionisation, operated as previously described.(Werner et al. 2019)

**Table 1:** Composition of mobile phase and elution gradient for targeted lipidomics.

| Time [min] | Flow rate [mL/min] | Eluent A [%]<br>(H<sub>2</sub>O/MeOH 90/10 +<br>0.01 % CH<sub>3</sub>COOH) | Eluent B [%]<br>(MeOH + 0.01 %<br>CH<sub>3</sub>COOH) |
| --- | --- | --- | --- |
| 0 | 0.3 | 64.4 | 35.6 |
| 4.8 - 19.0 | 0.3 | 45.5 | 54.5 |
| 26.7 | 0.3 | 15.6 | 84.4 |
| 29.7 | 0.3 | 2.2 | 97.8 |
| 30.7 | 0.3 | 64.4 | 35.6 |

The QTRAP 5500 was operated in negative mode using scheduled multiple reaction monitoring (MRM) with the MRM window set to 60 s. The retention time and constitutional stereochemistry of analytes were confirmed by external standards (Cayman Chemical), and individual calibration curves were obtained for quantification, using the ratios of the areas of external standards to their corresponding internal standard (ES/IS). Additional information, such as the specific transitions, detection and quantitation limits for each screened metabolite, is listed in Table S1. Representative unsmoothed chromatograms from samples showing peaks of oxylipins relevant to this study are provided in Fig. S4. For further analysis, the determined concentration was normalised to pg or ng oxylipin per 50 mg organ. The normalised raw data is provided in Table S2.

### Assessment of serum proteome (Proteome Profiler)

Serum samples were pooled from 4-5 biological replicates to a final volume of 100 µL. The serum samples were assessed for the relative abundance of 111 cytokines using the commercially available Proteome Profiler Mouse XL Cytokine Array Kit (ARY028) by R&D Systems, following the manufacturer’s guidelines. After treatment with IRDye 800CW streptavidin (LI-COR, Bad Homburg, Germany), and the membranes were analysed with an LI-COR Odyssey Infrared Imaging System. Pictures of the raw blots are included in Fig. S1.

### Data handling and statistical analysis

Results are presented as mean ± standard error of mean (SEM) unless stated otherwise. For data analysis and visualisation GraphPad Prism (Version 10.4.1), Origin Pro (Version 2023b), RStudio (2024.12.0) and Adobe Illustrator (version 29.3) were used. For the analysis of the oxylipin data, values below the determined limits of detection (LOD) were set to the corresponding LOD, and values between LOD and the lower limit of quantification (LLOQ) were set to ½ LLOQ, unless stated otherwise. For the principal component analysis, RStudio in combination with the R packages FactoMineR (https://cran.r-project.org/web/packages/FactoMineR/index.html) and factoextra (https://cran.r-project.org/web/packages/factoextra/index.html) was used, and metabolites for which all values were below the LLOQ were excluded from analysis. The ROUT outlier test was used to identify outliers within data sets, with a *Q*-value set to 0.5. Detected outliers were excluded from further analysis. The distribution of datasets was checked by Shapiro-Wilk test with α set to 0.05 and, if implied, data was log-transformed for statistical analysis. For group comparisons, an unpaired, two-tailed Student’s *t*-test was performed. Statistical significance was assumed for comparisons with *p* ≤ 0.05. Significance is indicated as: \**p* ≤ 0.05, \*\**p* ≤ 0.01, \*\*\**p* ≤ 0.001, \*\*\*\**p* ≤ 0.0001, or # respectively, ns, not significant.

## Declarations

### Ethical Statement

All studies were conducted in compliance with the recommendations of the European Commission for the protection of animals used for scientific purposes and with the approval of the local authorities Thüringer Landesamt für Verbraucherschutz, Germany license UKJ-19-014, and TWZ09-2022.

### Consent for publication

Not applicable.

### Availability of data and materials

The datasets analysed during the current study are included in this published article and its supplementary information files.

### CRediT author statement

**Mareike Wichmann-Costaganna:** Conceptualization, Methodology, Formal analysis, Investigation, Data Curation, Writing-Original Draft, Writing-Review & Editing, Visualization, Project Administration. **Raphaëlle Petit:** Methodology, Formal analysis, Investigation, Writing-Review & Editing. **Julia Lindner:** Methodology, Formal analysis, Investigation, Writing-Review & Editing. **Madlen Haase:** Methodology, Formal analysis, Investigation, Writing-Review & Editing. **Vivien Bachmann:** Investigation. **Robert Klaus Hofstetter:** Methodology, Validation, Writing-Review & Editing. **Markus Werner:** Methodology, Validation, Writing-Review & Editing. **Christiane Frahm:** Conceptualization, Supervision, Project administration, Resources, Writing-Review & Editing, Funding acquisition. **Oliver Werz:** Conceptualization, Resources, Writing-Original Draft, Writing-Review & Editing, Supervision, Project administration, Funding acquisition. **Patrick Schädel:** Conceptualization, Methodology, Formal analysis, Investigation, Data curation, Writing-Original Draft, Writing-Review & Editing, Supervision, Project administration.

## Acknowledgements

The authors acknowledge the kind support from the animal facility of UKJ and thank Katrin Fischer and Anna König for their excellent technical support. Schemes were created with biorender.com.

## Supplementary Information

**Figure S1:**
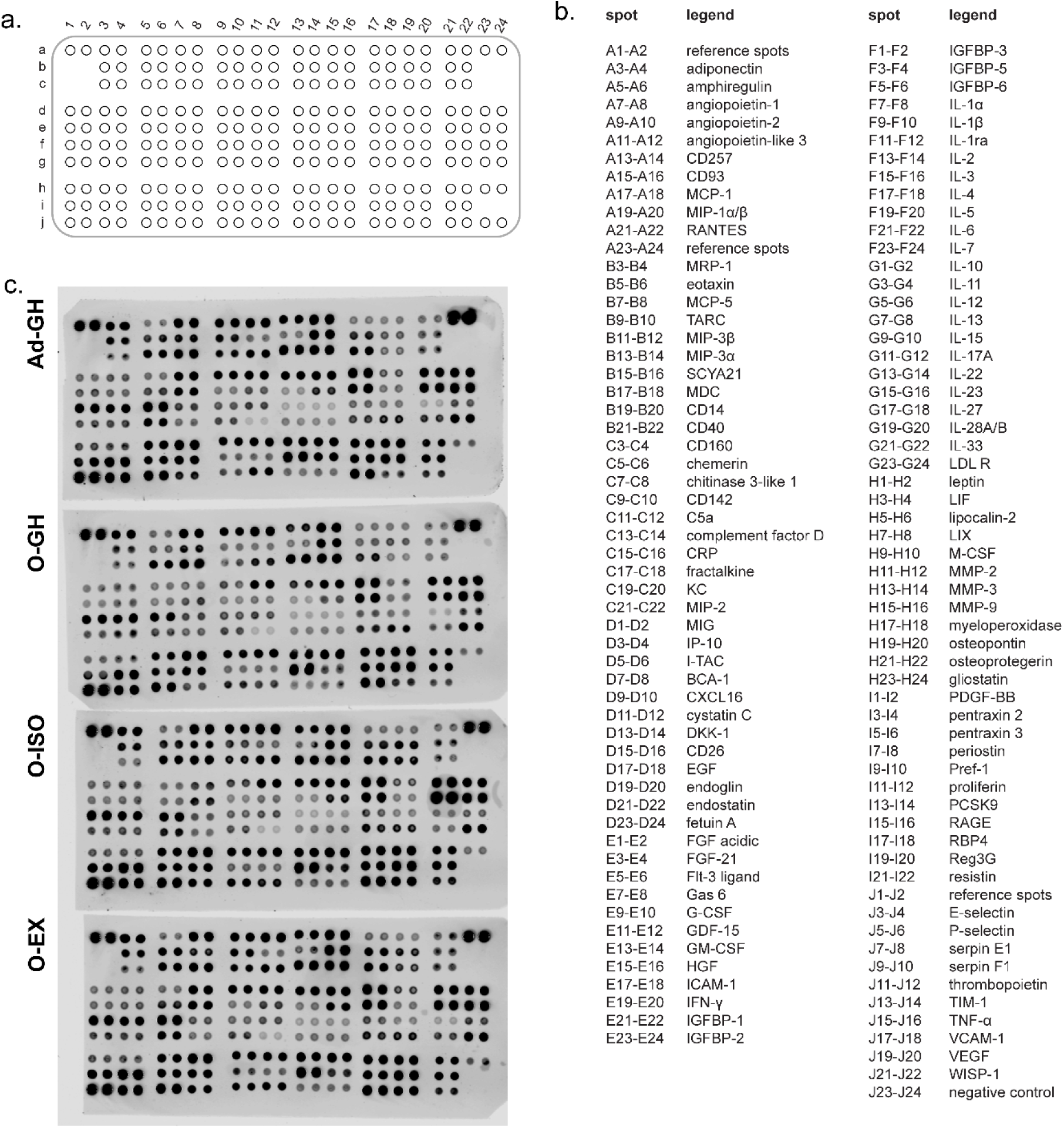
Raw pictures of proteome profiler membranes. **(a)** Schematic layout of the proteome profiler overlay template used for the analysis of circulating proteins in pooled serum samples. **(b)** List of the 111 screened proteins allotted to their corresponding position on the membrane. **(c)** Pictures of the membranes used for the analysis of circulating serum proteins in pooled serum samples.

**Figure S2:**
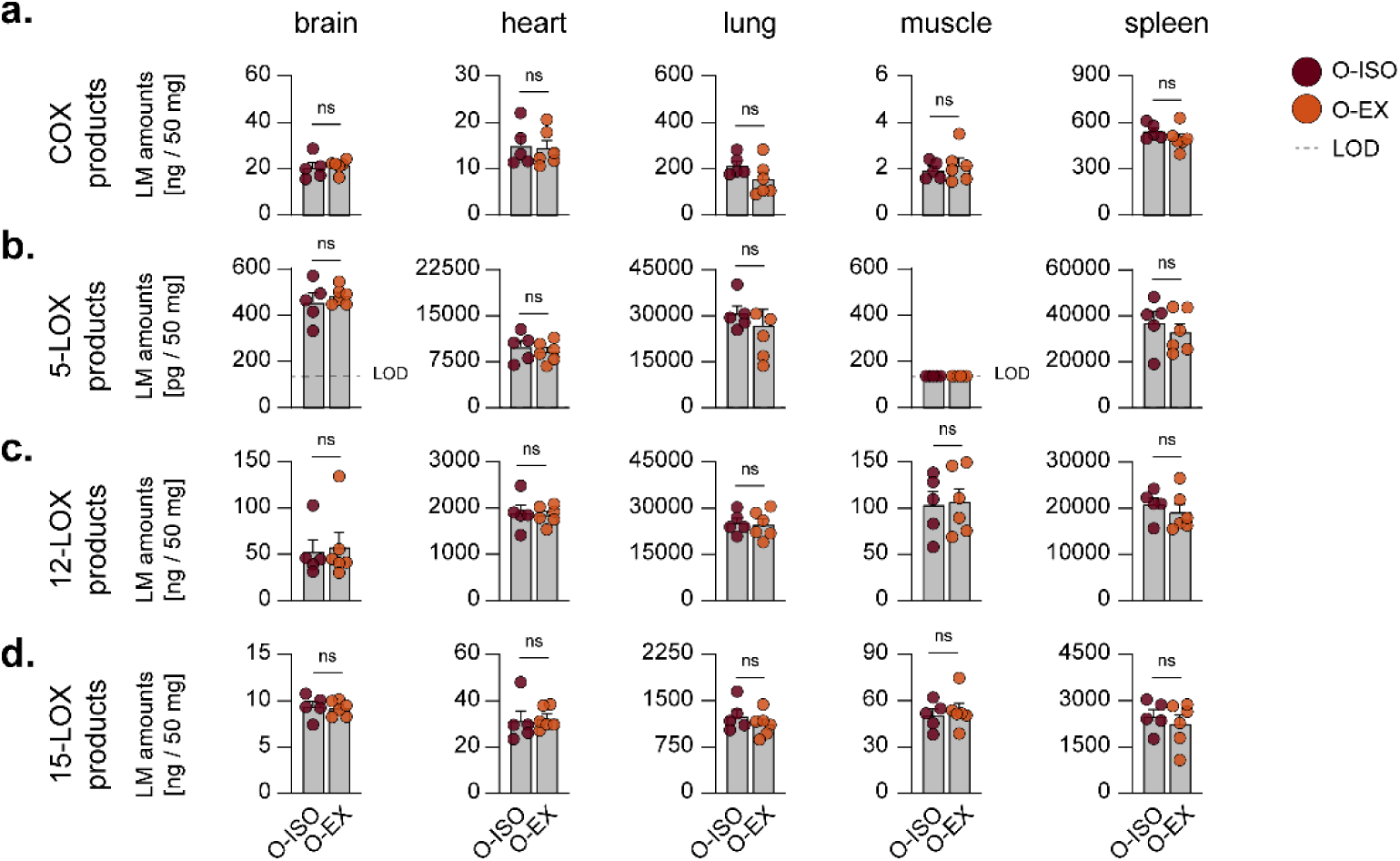
Enzyme-derived oxylipin levels for isolated mice with and without exercise. **(a-d)** Total amounts of grouped LM species for isolated mice with (O-EX) and without exercise (O-GH) that are products of the COX or LOX pathways in brain, heart, lung, muscle, spleen. Metabolites were grouped as follows: **(a)** *COX* – PGD_1_, PGE_1_, PGF_1α_, 6-keto PGF_1α_, PGD_2_, PGE_2_, 15-keto PGE_2_, PGF_2α_, PGF_2ß_, PGJ_2_, PGD_3_/PGE_3_, PGF_3α_, TXB_2_; **(b)** *5-LOX* – RvE1, RvE2, *trans*-LTB_4_, *epitrans*-LTB_4_, LTB_4_/5*S*,12*S*-diHETE, 5*S*,6*R*-diHETE, 20-OH LTB_4_, LTB_5_, 5-HETE, 5-HEPE, 7-HDHA; **(c)** *12-LOX* – MaR1, MaR2, 12-HETE, 12-HEPE, 14-HDHA; **(d)** *15-LOX* – PDx, PD1, RvD1, RvD2, RvD3, RvD4, RvD5, RvE4, LXA4, LXB4, LXA5, 5*S*,15*S*-diHETE, 15-HETE, 15-HEPE, 17-HDHA. Values are given as ng per 50 mg organ. LOD of the metabolites is indicated, if applicable.

**Figure S3:**
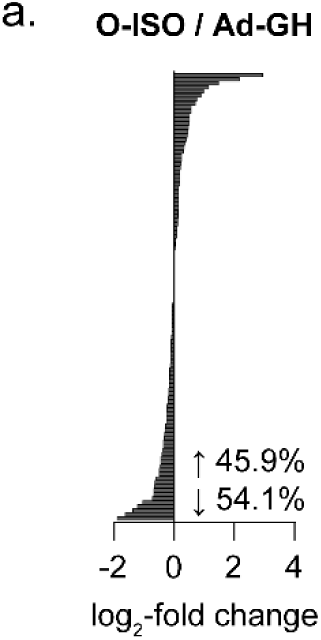
Circulatory proteome changes. **(a)** Log_2_-fold changes of 111 screened, circulating proteins in pooled serum samples and percentages of up- or downregulated proteins for the comparison of aged, isolated mice (O-ISO) against adult, group-housed (Ad-GH) mice.

**Figure S4:**
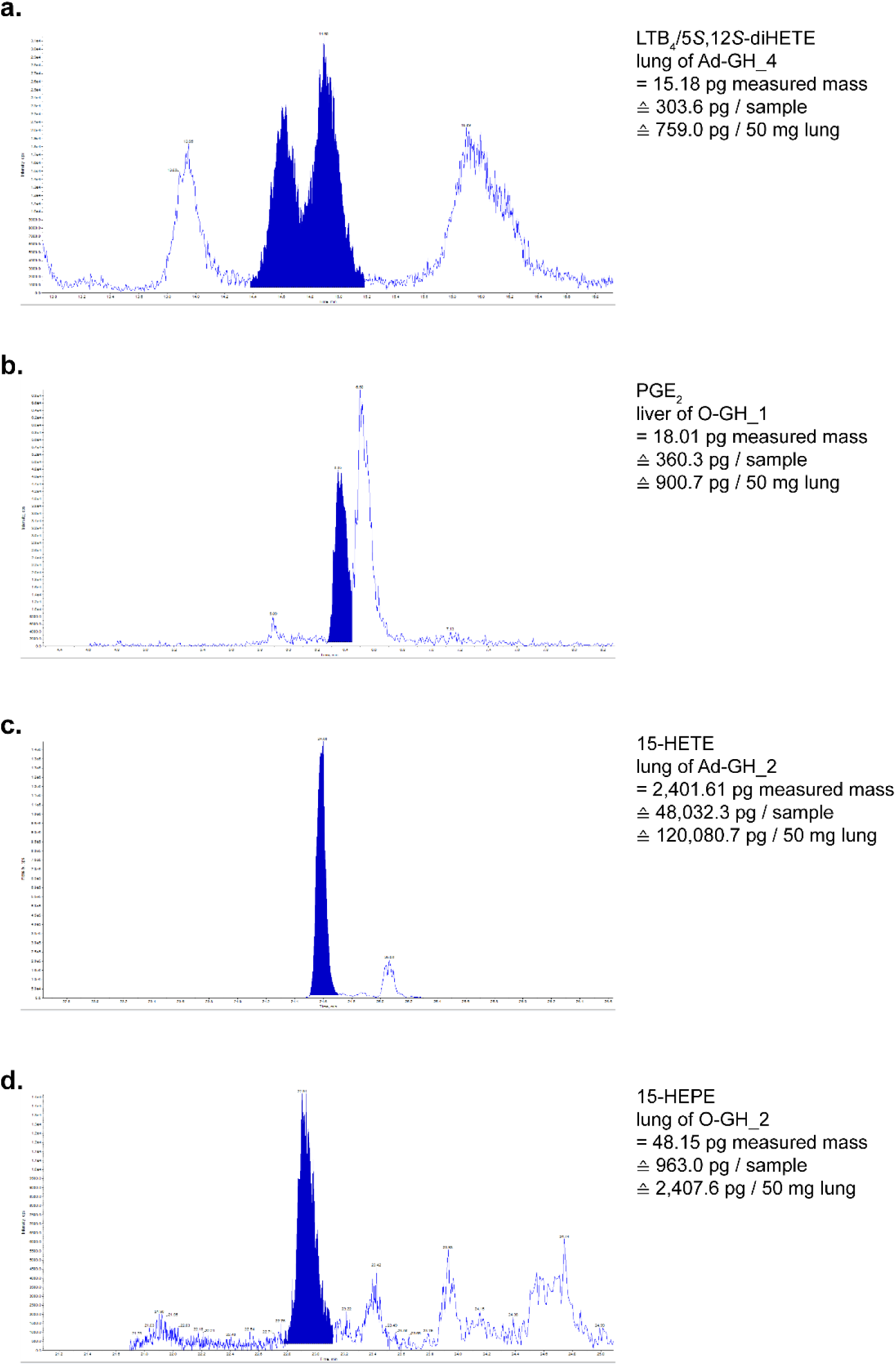
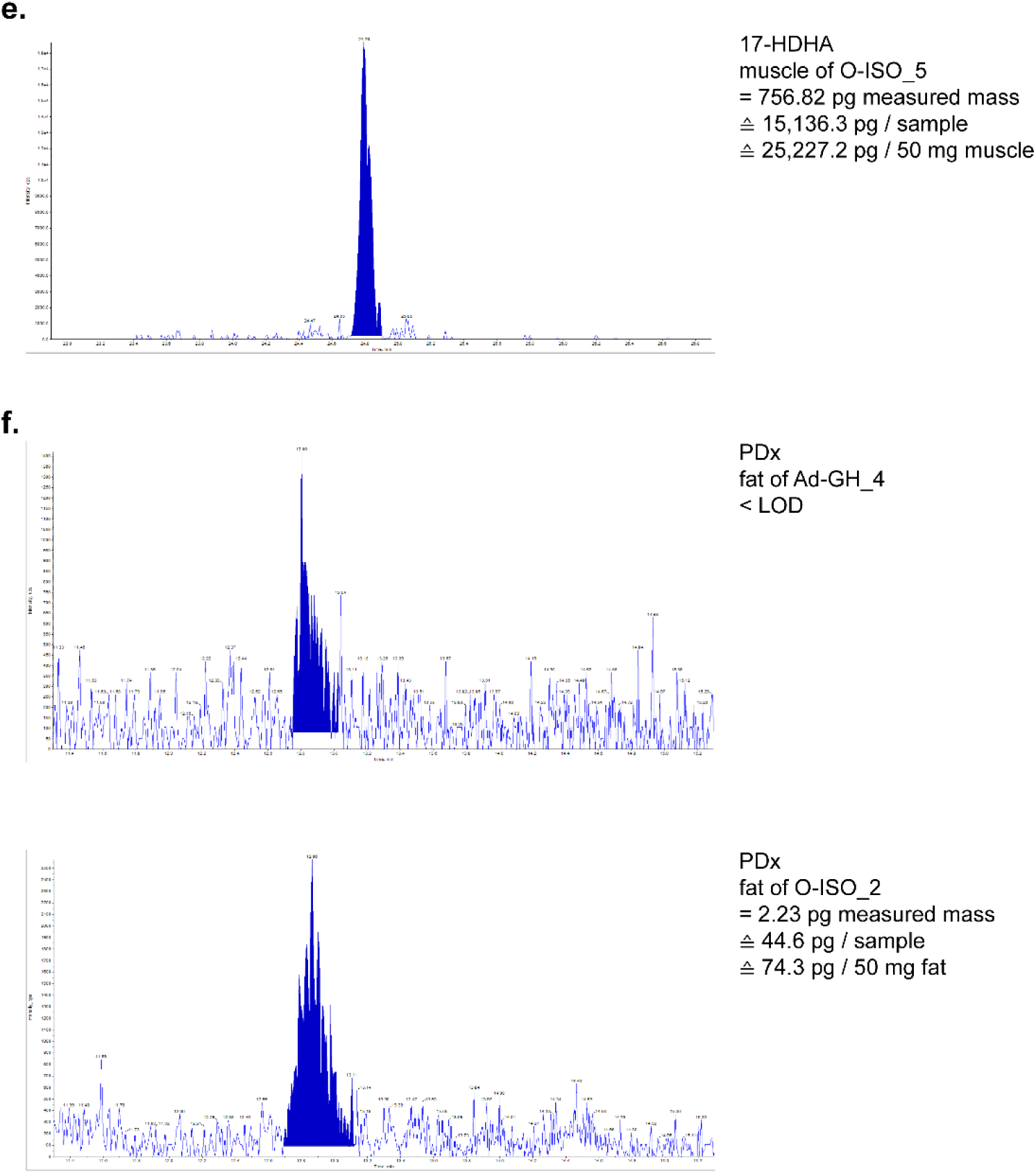
Representative chromatograms. Representative. unsmoothed chromatograms including the integration of relevant metabolites **(a)** LTB_4_/5*S*,12*S*-diHETE, **(b)** PGE_2_, **(c)** 15-HETE, **(d)** 15-HEPE, **(e)** 17-HDHA, and **(f)** PDx. Given values represent the measured mass on column, the calculated mass in the sample, and the normalized amount per 50 mg organ.

***Table S1: Screened metabolites, RT, transitions, LOD and LLOQ***

***Table S2: Oxylipin raw data***

## List of abbreviations

AA: arachidonic acid
Ad: adult
APP: acute-phase proteins
CD: cluster of differentiation
COX: cyclooxygenase
CRP: C-reactive protein
DHA: docosahexaenoic acid
EPA: eicosatetraenoic acid
EX: exercise
GC: glucocorticoid
GH: group-housed
HDHA: hydroxydocosahexaenoic acid
HETE: hydroxyeicosatetraenoic acid
HPA: hypothalamic-pituitary-adrenal
IL: Interleukin
IL-1ra: IL-1 receptor antagonist
ISO: isolated for 3 individual nights per week
LLOQ: limit of lower quantification
LOD: limit of detection
LOX: lipoxygenase
LT: leukotriene
LX: lipoxin
MaR: maresin
MRM: multiple reaction monitoring
O: aged
PG: prostaglandin
Rv: resolvin
SEM: standard error of mean
UPLC-MS/MS: ultra-performance liquid chromatography-tandem mass spectrometry
VEGF: vascular endothelial growth factor

## References

Akiyama, S., H. Nagai, S. Oike, I. Horikawa, M. Shinohara, Y. Lu, T. Futamura, R. Shinohara, S. Kitaoka, and T. Furuyashiki. 2022. ‘Chronic social defeat stress increases the amounts of 12-lipoxygenase lipid metabolites in the nucleus accumbens of stress-resilient mice’, Sci Rep, 12: 11385.

Arnardottir, Hildur H., Jesmond Dalli, Romain A. Colas, Masakazu Shinohara, and Charles N. Serhan. 2014. ‘Aging Delays Resolution of Acute Inflammation in Mice: Reprogramming the Host Response with Novel Nano-Proresolving Medicines’, The Journal of Immunology, 193: 4235–44.

Bäck, Magnus, and Göran K. Hansson. 2019. ‘Omega-3 fatty acids, cardiovascular risk, and the resolution of inflammation’, The FASEB Journal, 33: 1536–39.

Bender, G., E. E. Schexnaydre, R. C. Murphy, C. Uhlson, and M. E. Newcomer. 2016. ‘Membrane-dependent Activities of Human 15-LOX-2 and Its Murine Counterpart: IMPLICATIONS FOR MURINE MODELS OF ATHEROSCLEROSIS’, J Biol Chem, 291: 19413–24.

Berry, A., V. Bellisario, S. Capoccia, P. Tirassa, A. Calza, E. Alleva, and F. Cirulli. 2012. ‘Social deprivation stress is a triggering factor for the emergence of anxiety- and depression-like behaviours and leads to reduced brain BDNF levels in C57BL/6J mice’, Psychoneuroendocrinology, 37: 762–72.

Calderin, E. P., J. J. Zheng, N. L. Boyd, L. McNally, T. N. Audam, P. Lorkiewicz, B. G. Hill, and J. Hellmann. 2022. ‘Exercise-induced specialized proresolving mediators stimulate AMPK phosphorylation to promote mitochondrial respiration in macrophages’, Mol Metab, 66: 101637.

Cardona, M., and P. Andres. 2023. ‘Are social isolation and loneliness associated with cognitive decline in ageing?’, Front Aging Neurosci, 15: 1075563.

Chiurchiu, V., A. Leuti, and M. Maccarrone. 2018. ‘Bioactive Lipids and Chronic Inflammation: Managing the Fire Within’, Front Immunol, 9: 38.

Chiurchiu, V., M. Tiberi, A. Matteocci, F. Fazio, H. Siffeti, S. Saracini, N. B. Mercuri, and G. Sancesario. 2022. ’Lipidomics of Bioactive Lipids in Alzheimer’s and Parkinson’s Diseases: Where Are We?’, Int J Mol Sci, 23.

Cole, S. W., L. C. Hawkley, J. M. Arevalo, C. Y. Sung, R. M. Rose, and J. T. Cacioppo. 2007. ‘Social regulation of gene expression in human leukocytes’, Genome Biol, 8: R189.

Durham, W. J., S. L. Casperson, E. L. Dillon, M. A. Keske, D. Paddon-Jones, A. P. Sanford, R. C. Hickner, J. J. Grady, and M. Sheffield-Moore. 2010. ‘Age-related anabolic resistance after endurance-type exercise in healthy humans’, FASEB J, 24: 4117–27.

Ederer, M. L., M. Gunther, L. Best, J. Lindner, C. Kaleta, O. W. Witte, R. Simon, and C. Frahm. 2022. ‘Voluntary Wheel Running in Old C57BL/6 Mice Reduces Age-Related Inflammation in the Colon but Not in the Brain’, Cells, 11.

Ehrchen, J., L. Steinmuller, K. Barczyk, K. Tenbrock, W. Nacken, M. Eisenacher, U. Nordhues, C. Sorg, C. Sunderkotter, and J. Roth. 2007. ‘Glucocorticoids induce differentiation of a specifically activated, anti-inflammatory subtype of human monocytes’, Blood, 109: 1265–74.

Fenton, S. A. M., Jjcs Veldhuijzen van Zanten, J. L. Duda, G. S. Metsios, and G. D. Kitas. 2018. ‘Sedentary behaviour in rheumatoid arthritis: definition, measurement and implications for health’, Rheumatology (Oxford), 57: 213–26.

Ferrucci, L., and E. Fabbri. 2018. ‘Inflammageing: chronic inflammation in ageing, cardiovascular disease, and frailty’, Nat Rev Cardiol, 15: 505–22.

Fitzgerald, K. N., R. Hodges, D. Hanes, E. Stack, D. Cheishvili, M. Szyf, J. Henkel, M. W. Twedt, D. Giannopoulou, J. Herdell, S. Logan, and R. Bradley. 2021. ‘Potential reversal of epigenetic age using a diet and lifestyle intervention: a pilot randomized clinical trial’, Aging (Albany NY), 13: 9419–32.

Franceschi, C., M. Bonafe, S. Valensin, F. Olivieri, M. De Luca, E. Ottaviani, and G. De Benedictis. 2000. ‘Inflamm-aging. An evolutionary perspective on immunosenescence’, Ann N Y Acad Sci, 908: 244–54.

Franceschi, C., P. Garagnani, P. Parini, C. Giuliani, and A. Santoro. 2018. ‘Inflammaging: a new immune-metabolic viewpoint for age-related diseases’, Nat Rev Endocrinol, 14: 576–90.

Fulop, T., A. Larbi, G. Pawelec, A. Khalil, A. A. Cohen, K. Hirokawa, J. M. Witkowski, and C. Franceschi. 2023. ‘Immunology of Aging: the Birth of Inflammaging’, Clin Rev Allergy Immunol, 64: 109–22.

Funk, C. D. 2001. ‘Prostaglandins and leukotrienes: advances in eicosanoid biology’, Science, 294: 1871–5.

Furman, D., J. Campisi, E. Verdin, P. Carrera-Bastos, S. Targ, C. Franceschi, L. Ferrucci, D. W. Gilroy, A. Fasano, G. W. Miller, A. H. Miller, A. Mantovani, C. M. Weyand, N. Barzilai, J. J. Goronzy, T. A. Rando, R. B. Effros, A. Lucia, N. Kleinstreuer, and G. M. Slavich. 2019. ‘Chronic inflammation in the etiology of disease across the life span’, Nat Med, 25: 1822–32.

Garrido, A., I. Martinez de Toda, E. Diaz Del Cerro, J. Felix, N. Ceprian, M. Gonzalez-Sanchez, and M. De la Fuente. 2022. ‘Social environment as a modulator of immunosenescence’, Expert Rev Mol Med, 24: e29.

Gleeson, M., N. C. Bishop, D. J. Stensel, M. R. Lindley, S. S. Mastana, and M. A. Nimmo. 2011. ‘The anti-inflammatory effects of exercise: mechanisms and implications for the prevention and treatment of disease’, Nat Rev Immunol, 11: 607–15.

Gleim, S., J. Stitham, W. H. Tang, K. A. Martin, and J. Hwa. 2012. ‘An eicosanoid-centric view of atherothrombotic risk factors’, Cell Mol Life Sci, 69: 3361–80.

Goppelt-Struebe, M., D. Wolter, and K. Resch. 1989. ‘Glucocorticoids inhibit prostaglandin synthesis not only at the level of phospholipase A2 but also at the level of cyclo-oxygenase/PGE isomerase’, Br J Pharmacol, 98: 1287–95.

Grant, N., M. Hamer, and A. Steptoe. 2009. ‘Social isolation and stress-related cardiovascular, lipid, and cortisol responses’, Ann Behav Med, 37: 29–37.

Gutierrez, F., M. Masia, C. Mirete, B. Soldan, J. C. Rodriguez, S. Padilla, I. Hernandez, G. Royo, and A. Martin-Hidalgo. 2006. ‘The influence of age and gender on the population-based incidence of community-acquired pneumonia caused by different microbial pathogens’, J Infect, 53: 166–74.

Haeggstrom, J. Z. 2018. ‘Leukotriene biosynthetic enzymes as therapeutic targets’, J Clin Invest, 128: 2680–90.

Hammig, O. 2019. ‘Health risks associated with social isolation in general and in young, middle and old age’, PLoS One, 14: e0219663.

Harman, D. 2006. ‘Free radical theory of aging: an update: increasing the functional life span’, Ann N Y Acad Sci, 1067: 10–21.

Hodgson, S., I. Watts, S. Fraser, P. Roderick, and H. Dambha-Miller. 2020. ‘Loneliness, social isolation, cardiovascular disease and mortality: a synthesis of the literature and conceptual framework’, J R Soc Med, 113: 185–92.

Holt-Lunstad, J., T. B. Smith, and J. B. Layton. 2010. ‘Social relationships and mortality risk: a meta-analytic review’, PLoS Med, 7: e1000316.

Khan, S. S., B. D. Singer, and D. E. Vaughan. 2017. ‘Molecular and physiological manifestations and measurement of aging in humans’, Aging Cell, 16: 624–33.

Larsson, S. C., J. Kaluza, and A. Wolk. 2017. ‘Combined impact of healthy lifestyle factors on lifespan: two prospective cohorts’, J Intern Med, 282: 209–19.

Leigh-Hunt, N., D. Bagguley, K. Bash, V. Turner, S. Turnbull, N. Valtorta, and W. Caan. 2017. ‘An overview of systematic reviews on the public health consequences of social isolation and loneliness’, Public Health, 152: 157–71.

Leuti, A., D. Fazio, M. Fava, A. Piccoli, S. Oddi, and M. Maccarrone. 2020. ‘Bioactive lipids, inflammation and chronic diseases’, Adv Drug Deliv Rev, 159: 133–69.

Levy, B. D., C. B. Clish, B. Schmidt, K. Gronert, and C. N. Serhan. 2001. ‘Lipid mediator class switching during acute inflammation: signals in resolution’, Nat Immunol, 2: 612–9.

Lopez-Otin, C., M. A. Blasco, L. Partridge, M. Serrano, and G. Kroemer. 2023. ‘Hallmarks of aging: An expanding universe’, Cell, 186: 243–78.

Lopez-Otin, C., L. Galluzzi, J. M. P. Freije, F. Madeo, and G. Kroemer. 2016. ‘Metabolic Control of Longevity’, Cell, 166: 802–21.

Luo, Y., S. Kuang, L. Xue, and J. Yang. 2016. ‘The mechanism of 5-lipoxygenase in the impairment of learning and memory in rats subjected to chronic unpredictable mild stress’, Physiol Behav, 167: 145–53.

Magalhaes, D. M., M. Mampay, A. M. Sebastiao, G. K. Sheridan, and C. A. Valente. 2024. ‘Age-related impact of social isolation in mice: Young vs middle-aged’, Neurochem Int, 174: 105678.

Malan, L., L. Zandberg, C. Pienaar, A. Nienaber, and L. Havemann-Nel. 2024. ‘Regular moderate physical activity potentially accelerates and strengthens both the pro-inflammatory and pro-resolving lipid mediator response after acute exercise stress’, Prostaglandins Leukot Essent Fatty Acids, 202: 102642.

Markworth, J. F., K. R. Maddipati, and D. Cameron-Smith. 2016. ‘Emerging roles of pro-resolving lipid mediators in immunological and adaptive responses to exercise-induced muscle injury’.

McKim, D. B., W. Yin, Y. Wang, S. W. Cole, J. P. Godbout, and J. F. Sheridan. 2018. ‘Social Stress Mobilizes Hematopoietic Stem Cells to Establish Persistent Splenic Myelopoiesis’, Cell Rep, 25: 2552–62 e3.

Minhas, P. S., A. Latif-Hernandez, M. R. McReynolds, A. S. Durairaj, Q. Wang, A. Rubin, A. U. Joshi, J. Q. He, E. Gauba, L. Liu, C. Wang, M. Linde, Y. Sugiura, P. K. Moon, R. Majeti, M. Suematsu, D. Mochly-Rosen, I. L. Weissman, F. M. Longo, J. D. Rabinowitz, and K. I. Andreasson. 2021. ‘Restoring metabolism of myeloid cells reverses cognitive decline in ageing’, Nature, 590: 122–28.

Mrowetz, H., M. H. Kotob, J. Forster, I. Aydin, M. S. Unger, J. Lubec, A. M. Hussein, J. Malikovic, D. D. Feyissa, V. Korz, H. Hoger, G. Lubec, and L. Aigner. 2023. ‘Leukotriene signaling as molecular correlate for cognitive heterogeneity in aging: an exploratory study’, Front Aging Neurosci, 15: 1140708.

Mumtaz, F., M. I. Khan, M. Zubair, and A. R. Dehpour. 2018. ‘Neurobiology and consequences of social isolation stress in animal model-A comprehensive review’, Biomed Pharmacother, 105: 1205–22.

Muta, O., M. Odaka, Y. Fujii, T. Fushimi, H. Sato, and N. Osakabe. 2023. ‘Difference in endocrine and behavior between short-term single- and paired-housing mice in metabolic cage’, Neurosci Lett, 806: 137246.

Nayeem, M. A. 2018. ‘Role of oxylipins in cardiovascular diseases’, Acta Pharmacol Sin, 39: 1142–54.

Ngandu, T., J. Lehtisalo, A. Solomon, E. Levalahti, S. Ahtiluoto, R. Antikainen, L. Backman, T. Hanninen, A. Jula, T. Laatikainen, J. Lindstrom, F. Mangialasche, T. Paajanen, S. Pajala, M. Peltonen, R. Rauramaa, A. Stigsdotter-Neely, T. Strandberg, J. Tuomilehto, H. Soininen, and M. Kivipelto. 2015. ‘A 2 year multidomain intervention of diet, exercise, cognitive training, and vascular risk monitoring versus control to prevent cognitive decline in at-risk elderly people (FINGER): a randomised controlled trial’, Lancet, 385: 2255–63.

Olecka, M., H. Morrison, and S. Hoffmann. 2025. ‘Signatures of Nonlinear Aging: Molecular Stages of Life: Sudden Changes During Aging as Potential Biomarkers for an Age Classification System’, Bioessays, 47: e202400222.

Olecka, M., A. van Bommel, L. Best, M. Haase, S. Foerste, K. Riege, T. Dost, S. Flor, O. W. Witte, S. Franzenburg, M. Groth, B. von Eyss, C. Kaleta, C. Frahm, and S. Hoffmann. 2024. ‘Nonlinear DNA methylation trajectories in aging male mice’, Nat Commun, 15: 3074.

Oliva, C. A., M. Lira, C. Jara, A. Catenaccio, T. A. Mariqueo, C. B. Lindsay, F. Bozinovic, G. Cavieres, N. C. Inestrosa, C. Tapia-Rojas, and D. S. Rivera. 2023. ’Long-term social isolation stress exacerbates sex-specific neurodegeneration markers in a natural model of Alzheimer’s disease’, Front Aging Neurosci, 15: 1250342.

Pels, Fabian, and Jens Kleinert. 2016. ‘Loneliness and physical activity: A systematic review’, International Review of Sport and Exercise Psychology, 9: 231–60.

Pena Calderin, E., J. J. Zheng, N. L. Boyd, W. Lynch, B. E. Sansbury, M. Spite, B. G. Hill, and J. Hellmann. 2025. ‘Exercise-Stimulated Resolvin Biosynthesis in the Adipose Tissue Is Abrogated by High-Fat Diet-Induced Adrenergic Deficiency’, Arterioscler Thromb Vasc Biol, 45: 1090–110.

Rao, Z., E. Brunner, B. Giszas, A. Iyer-Bierhoff, J. Gerstmeier, F. Borner, P. M. Jordan, S. Pace, K. P. L. Meyer, R. K. Hofstetter, D. Merk, C. Paulenz, T. Heinzel, P. C. Grunert, A. Stallmach, C. N. Serhan, M. Werner, and O. Werz. 2023. ‘Glucocorticoids regulate lipid mediator networks by reciprocal modulation of 15-lipoxygenase isoforms affecting inflammation resolution’, Proc Natl Acad Sci U S A, 120: e2302070120.

Riddick, C. A., W. L. Ring, J. R. Baker, C. R. Hodulik, and T. D. Bigby. 1997. ‘Dexamethasone increases expression of 5-lipoxygenase and its activating protein in human monocytes and THP-1 cells’, Eur J Biochem, 246: 112–8.

Ruiz, L. A., R. Zalacain, A. Capelastegui, A. Bilbao, A. Gomez, A. Uranga, and P. P. Espana. 2014. ‘Bacteremic pneumococcal pneumonia in elderly and very elderly patients: host-and pathogen-related factors, process of care, and outcome’, J Gerontol A Biol Sci Med Sci, 69: 1018–24.

Schadel, P., M. Wichmann-Costaganna, A. Czapka, N. Gebert, A. Ori, and O. Werz. 2023. ‘Short-Term Caloric Restriction and Subsequent Re-Feeding Compromise Liver Health and Associated Lipid Mediator Signaling in Aged Mice’, Nutrients, 15.

Schädel, Patrick, Fabiana Troisi, Anna Czapka, Nadja Gebert, Simona Pace, Alessandro Ori, and Oliver Werz. 2021. ‘Aging drives organ-specific alterations of the inflammatory microenvironment guided by immunomodulatory mediators in mice’, The FASEB Journal, 35.

Serhan, C. N. 2014. ‘Pro-resolving lipid mediators are leads for resolution physiology’, Nature, 510: 92–101.

Shen, X., C. Wang, X. Zhou, W. Zhou, D. Hornburg, S. Wu, and M. P. Snyder. 2024. ‘Nonlinear dynamics of multi-omics profiles during human aging’, Nat Aging.

Shoji, H., and K. Mizoguchi. 2011. ‘Aging-related changes in the effects of social isolation on social behavior in rats’, Physiol Behav, 102: 58–62.

Smith, K. J., S. Gavey, R. Iddell NE, P. Kontari, and C. Victor. 2020. ‘The association between loneliness, social isolation and inflammation: A systematic review and meta-analysis’, Neurosci Biobehav Rev, 112: 519–41.

Smolensky, I., K. Zajac-Bakri, A. S. Mallien, P. Gass, R. Guzman, and D. Inta. 2024. ‘Effects of single housing on behavior, corticosterone level and body weight in male and female mice’, Lab Anim Res, 40: 35.

Sun, S., S. Ma, Y. Cai, S. Wang, J. Ren, Y. Yang, J. Ping, X. Wang, Y. Zhang, H. Yan, W. Li, C. R. Esteban, Y. Yu, F. Liu, J. C. Izpisua Belmonte, W. Zhang, J. Qu, and G. H. Liu. 2023. ‘A single-cell transcriptomic atlas of exercise-induced anti-inflammatory and geroprotective effects across the body’, Innovation (Camb), 4: 100380.

Umamaheswaran, S., S. K. Dasari, P. Yang, S. K. Lutgendorf, and A. K. Sood. 2018. ‘Stress, inflammation, and eicosanoids: an emerging perspective’, Cancer Metastasis Rev, 37: 203–11.

Wang, F., Y. Gao, Z. Han, Y. Yu, Z. Long, X. Jiang, Y. Wu, B. Pei, Y. Cao, J. Ye, M. Wang, and Y. Zhao. 2023. ‘A systematic review and meta-analysis of 90 cohort studies of social isolation, loneliness and mortality’, Nat Hum Behav, 7: 1307–19.

Wang, J., C. Chen, J. Zhou, L. Ye, Y. Li, L. Xu, Z. Xu, X. Li, Y. Wei, J. Liu, Y. Lv, and X. Shi. 2023. ‘Healthy lifestyle in late-life, longevity genes, and life expectancy among older adults: a 20-year, population-based, prospective cohort study’, Lancet Healthy Longev, 4: e535–e43.

Werner, Markus, Paul M. Jordan, Erik Romp, Anna Czapka, Zhigang Rao, Christian Kretzer, Andreas Koeberle, Ulrike Garscha, Simona Pace, Hans-Erik Claesson, Charles N. Serhan, Oliver Werz, and Jana Gerstmeier. 2019. ‘Targeting biosynthetic networks of the proinflammatory and proresolving lipid metabolome’, The FASEB Journal, 33: 6140–53.

Whittington, R. A., E. Planel, and N. Terrando. 2017. ’Impaired Resolution of Inflammation in Alzheimer’s Disease: A Review’, Front Immunol, 8: 1464.

Yang, Y. C., C. Boen, K. Gerken, T. Li, K. Schorpp, and K. M. Harris. 2016. ‘Social relationships and physiological determinants of longevity across the human life span’, Proc Natl Acad Sci U S A, 113: 578–83.

Yang, Y. C., M. K. McClintock, M. Kozloski, and T. Li. 2013. ‘Social isolation and adult mortality: the role of chronic inflammation and sex differences’, J Health Soc Behav, 54: 183–203.

Yi, B., M. Rykova, M. Feuerecker, B. Jager, C. Ladinig, M. Basner, M. Horl, S. Matzel, I. Kaufmann, C. Strewe, I. Nichiporuk, G. Vassilieva, K. Rinas, S. Baatout, G. Schelling, M. Thiel, D. F. Dinges, B. Morukov, and A. Chouker. 2014. ‘520-d Isolation and confinement simulating a flight to Mars reveals heightened immune responses and alterations of leukocyte phenotype’, Brain Behav Immun, 40: 203–10.

