## Supplementary Figures for "Social isolation of aged mice drives dramatic release of inflammatory lipoxygenase-derived oxylipins"

### Supplementary Information

### Figure S1: Raw pictures of proteome profiler membranes

**(a)** Schematic layout of the proteome profiler overlay template used for the analysis of circulating proteins in pooled serum samples. **(b)** List of the 111 screened proteins allotted to their corresponding position on the membrane. **(c)** Pictures of the membranes used for the analysis of circulating serum proteins in pooled serum samples.

a.

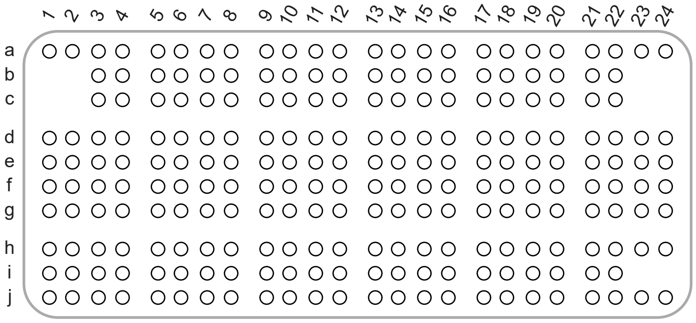


b.

| spot | legend | spot | legend |
|---|---|---|---|
| A1-A2 | reference spots | F1-F2 | IGFBP-3 |
| A3-A4 | adiponectin | F3-F4 | IGFBP-5 |
| A5-A6 | amphiregulin | F5-F6 | IGFBP-6 |
| A7-A8 | angiopoietin-1 | F7-F8 | IL-1α |
| A9-A10 | angiopoietin-2 | F9-F10 | IL-1β |
| A11-A12 | angiopoietin-like 3 | F11-F12 | IL-1ra |
| A13-A14 | CD257 | F13-F14 | IL-2 |
| A15-A16 | CD93 | F15-F16 | IL-3 |
| A17-A18 | MCP-1 | F17-F18 | IL-4 |
| A19-A20 | MIP-1α/β | F19-F20 | IL-5 |
| A21-A22 | RANTES | F21-F22 | IL-6 |
| A23-A24 | reference spots | F23-F24 | IL-7 |
| B3-B4 | MRP-1 | G1-G2 | IL-10 |
| B5-B6 | eotaxin | G3-G4 | IL-11 |
| B7-B8 | MCP-5 | G5-G6 | IL-12 |
| B9-B10 | TARC | G7-G8 | IL-13 |
| B11-B12 | MIP-3β | G9-G10 | IL-15 |
| B13-B14 | MIP-3α | G11-G12 | IL-17A |
| B15-B16 | SCYA21 | G13-G14 | IL-22 |
| B17-B18 | MDC | G15-G16 | IL-23 |
| B19-B20 | CD14 | G17-G18 | IL-27 |
| B21-B22 | CD40 | G19-G20 | IL-28A/B |
| C3-C4 | CD160 | G21-G22 | IL-33 |
| C5-C6 | chemerin | G23-G24 | LDL R |
| C7-C8 | chitinase 3-like 1 | H1-H2 | leptin |
| C9-C10 | CD142 | H3-H4 | LIF |
| C11-C12 | C5a | H5-H6 | lipocalin-2 |
| C13-C14 | complement factor D | H7-H8 | LIX |
| C15-C16 | CRP | H9-H10 | M-CSF |
| C17-C18 | fractalkine | H11-H12 | MMP-2 |
| C19-C20 | KC | H13-H14 | MMP-3 |
| C21-C22 | MIP-2 | H15-H16 | MMP-9 |
| D1-D2 | MIG | H17-H18 | myeloperoxidase |
| D3-D4 | IP-10 | H19-H20 | osteopontin |
| D5-D6 | I-TAC | H21-H22 | osteoprotegerin |
| D7-D8 | BCA-1 | H23-H24 | gliostatin |
| D9-D10 | CXCL16 | I1-I2 | PDGF-BB |
| D11-D12 | cystatin C | I3-I4 | pentraxin 2 |
| D13-D14 | DKK-1 | I5-I6 | pentraxin 3 |
| D15-D16 | CD26 | I7-I8 | periostin |
| D17-D18 | EGF | I9-I10 | Pref-1 |
| D19-D20 | endoglin | I11-I12 | proliferin |
| D21-D22 | endostatin | I13-I14 | PCSK9 |
| D23-D24 | fetuin A | I15-I16 | RAGE |
| E1-E2 | FGF acidic | I17-I18 | RBP4 |
| E3-E4 | FGF-21 | I19-I20 | Reg3G |
| E5-E6 | Flt-3 ligand | I21-I22 | resistin |
| E7-E8 | Gas 6 | J1-J2 | reference spots |
| E9-E10 | G-CSF | J3-J4 | E-selectin |
| E11-E12 | GDF-15 | J5-J6 | P-selectin |
| E13-E14 | GM-CSF | J7-J8 | serpin E1 |
| E15-E16 | HGF | J9-J10 | serpin F1 |
| E17-E18 | ICAM-1 | J11-J12 | thrombopoietin |
| E19-E20 | IFN-γ | J13-J14 | TIM-1 |
| E21-E22 | IGFBP-1 | J15-J16 | TNF-α |
| E23-E24 | IGFBP-2 | J17-J18 | VCAM-1 |
| | | J19-J20 | VEGF |
| | | J21-J22 | WISP-1 |
| | | J23-J24 | negative control |

c.

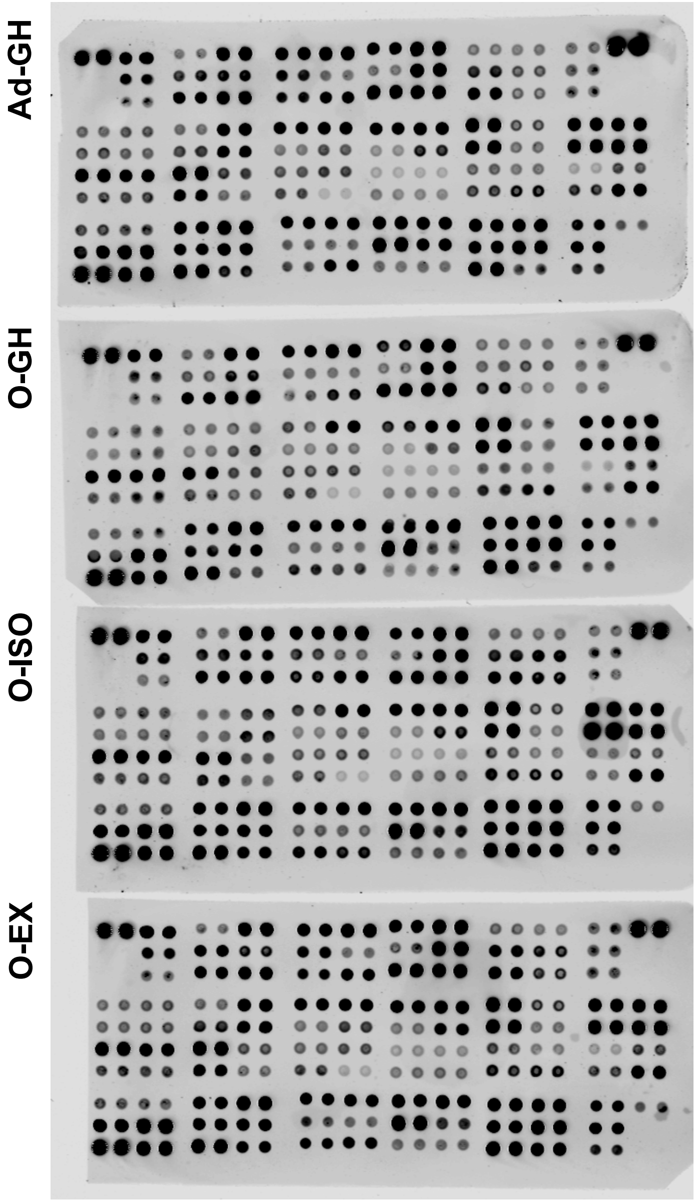

**Figure S2: Enzyme-derived oxylipin levels for isolated mice with and without exercise.**

**(a-d)** Total amounts of grouped LM species for isolated mice with (O-EX) and without exercise (O-GH) that are products of the COX or LOX pathways in brain, heart, lung, muscle, spleen. Metabolites were grouped as follows: **(a)** *COX* – $PGD_1$, $PGE_1$, $PGF_{1\alpha}$, 6-keto $PGF_{1\alpha}$, $PGD_2$, $PGE_2$, 15-keto $PGE_2$, $PGF_{2\alpha}$, $PGF_{2\beta}$, $PGJ_2$, $PGD_3$/$PGE_3$, $PGF_{3\alpha}$, $TXB_2$; **(b)** *5-LOX* – RvE1, RvE2, *trans*-$LTB_4$, *epitrans*-$LTB_4$, $LTB_4$/5*S*,12*S*-diHETE, 5*S*,6*R*-diHETE, 20-OH $LTB_4$, $LTB_5$, 5-HETE, 5-HEPE, 7-HDHA; **(c)** *12-LOX* – MaR1, MaR2, 12-HETE, 12-HEPE, 14-HDHA; **(d)** *15-LOX* –PDx, PD1, RvD1, RvD2, RvD3, RvD4, RvD5, RvE4, LXA4, LXB4, LXA5, 5*S*,15*S*-diHETE, 15-HETE, 15-HEPE, 17-HDHA. Values are given as ng per 50 mg organ. LOD of the metabolites is indicated, if applicable.

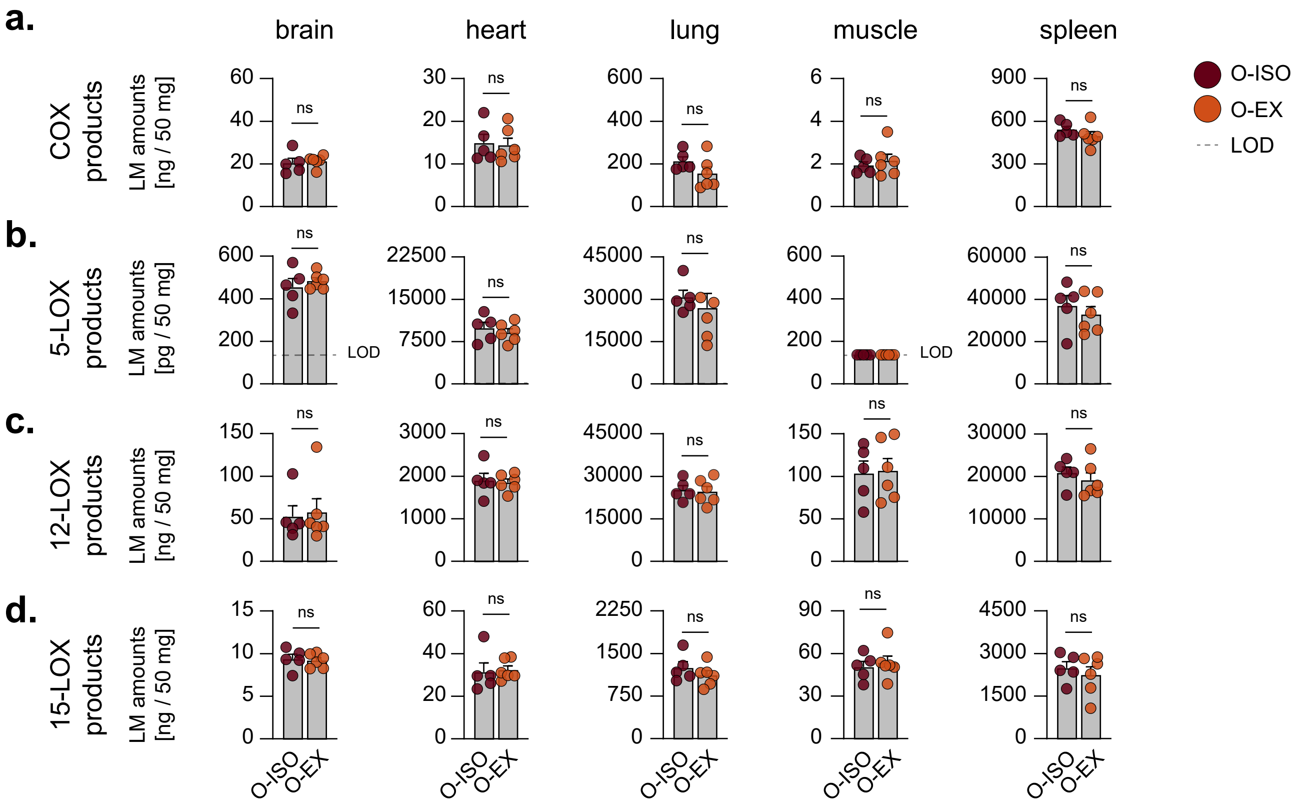

a.
b.
c.
d.
brain
heart
lung
muscle
spleen
COX products
5-LOX products
12-LOX products
15-LOX products
LM amounts [ng / 50 mg]
LM amounts [pg / 50 mg]
ns
LOD
O-ISO
O-EX

**Figure S3: Circulatory proteome changes.**

**(a)** $Log_2$-fold changes of 111 screened, circulating proteins in pooled serum samples and percentages of up- or downregulated proteins for the comparison of aged, isolated mice (O-ISO) against adult, group-housed (Ad-GH) mice.

a.

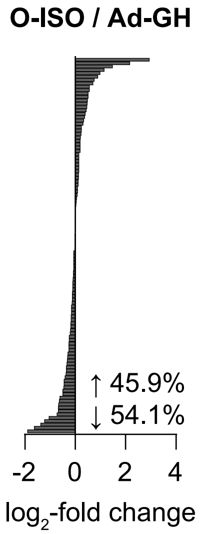

**Figure S4: Representative chromatograms.**

Representative. unsmoothed chromatograms including the integration of relevant metabolites **(a)** $LTB_4$/5*S*,12*S*-diHETE, **(b)** $PGE_2$, **(c)** 15-HETE, **(d)** 15-HEPE, **(e)** 17-HDHA, and **(f)** PDx. Given values represent the measured mass on column, the calculated mass in the sample, and the normalized amount per 50 mg organ.

**a.**

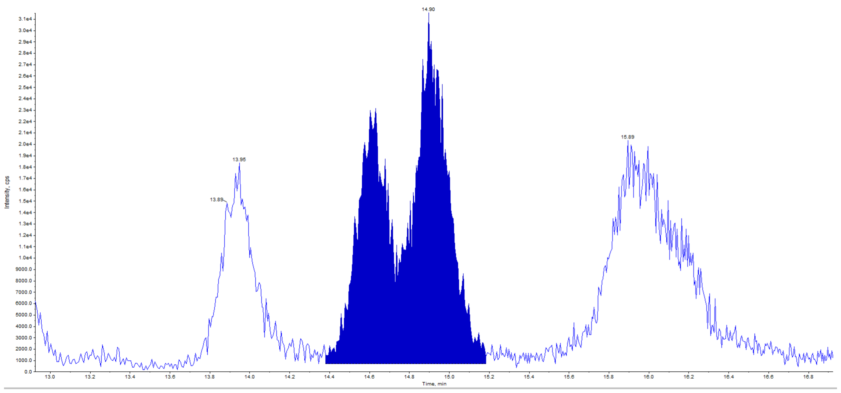

LTB$_4$/5*S*,12*S*-diHETE
lung of Ad-GH_4
= 15.18 pg measured mass
≙ 303.6 pg / sample
≙ 759.0 pg / 50 mg lung

**b.**

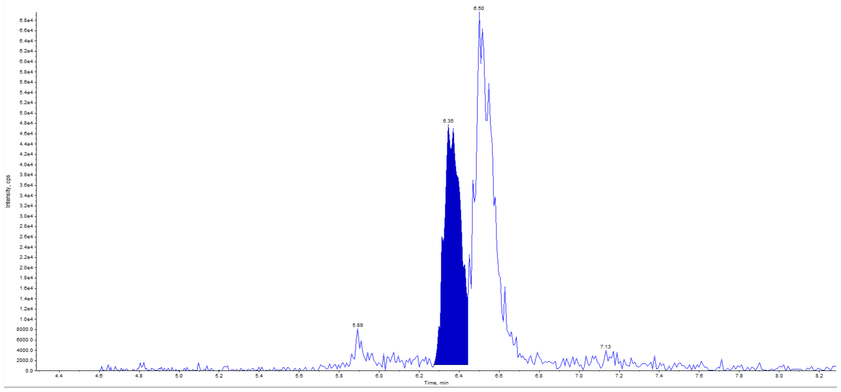

PGE$_2$
liver of O-GH_1
= 18.01 pg measured mass
≙ 360.3 pg / sample
≙ 900.7 pg / 50 mg lung

**c.**

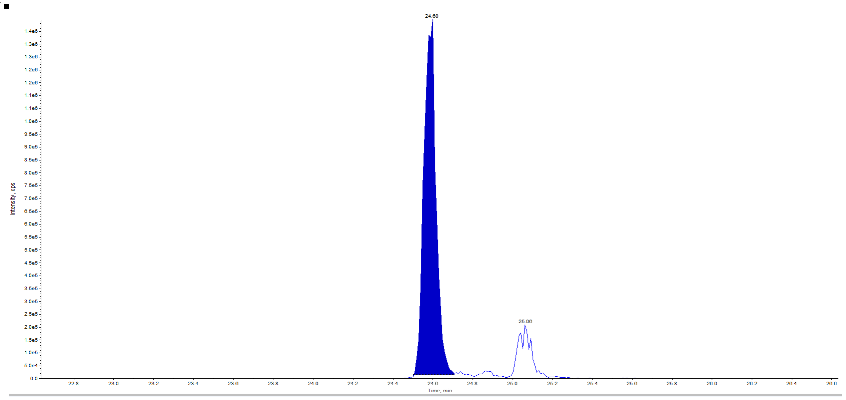

15-HETE
lung of Ad-GH_2
= 2,401.61 pg measured mass
≙ 48,032.3 pg / sample
≙ 120,080.7 pg / 50 mg lung

**d.**

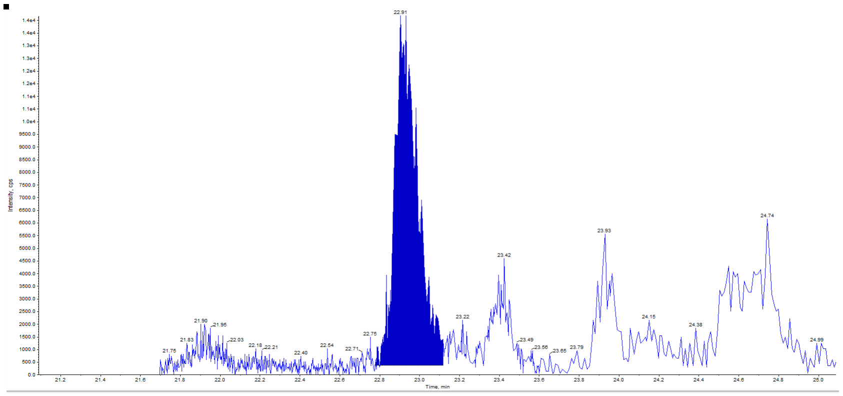

15-HEPE
lung of O-GH_2
= 48.15 pg measured mass
≙ 963.0 pg / sample
≙ 2,407.6 pg / 50 mg lung

**e.**

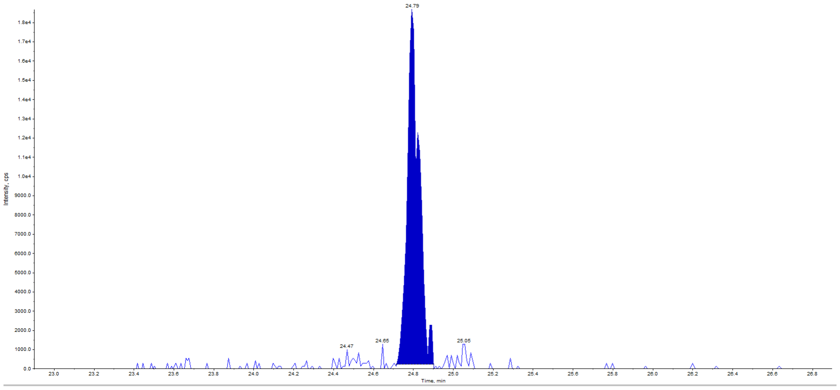

17-HDHA
muscle of O-ISO_5
= 756.82 pg measured mass
≙ 15,136.3 pg / sample
≙ 25,227.2 pg / 50 mg muscle

**f.**

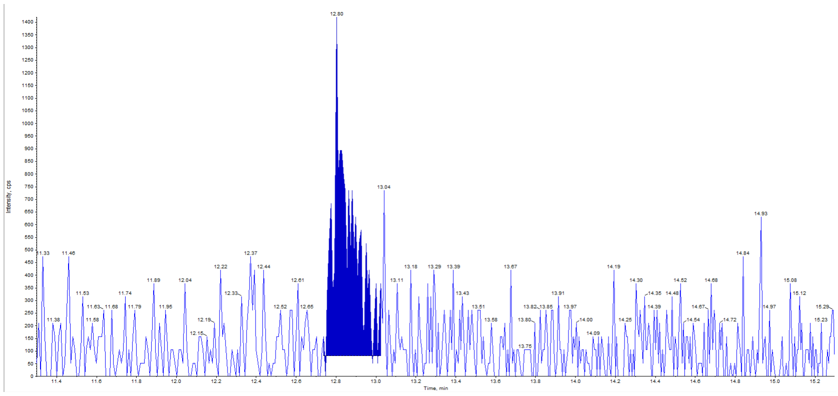

PDx
fat of Ad-GH_4
< LOD

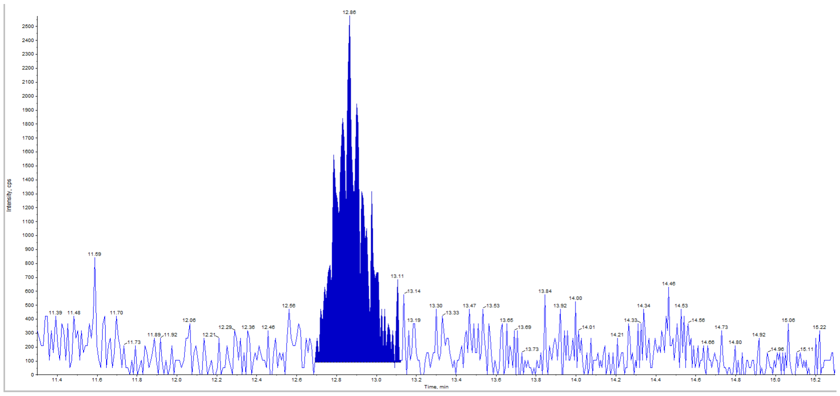

PDx
fat of O-ISO_2
= 2.23 pg measured mass
≙ 44.6 pg / sample
≙ 74.3 pg / 50 mg fat

**Table S1: Screened metabolites, transitions, LOD and LLOQ**

**Table S2: Oxylipin raw data**
